# Regulation of Hepatic MicroRNAs in Response to Early Stage *Echinococcus multilocularis* Egg Infection in C57BL/6 mice

**DOI:** 10.1101/708750

**Authors:** Ghalia Boubaker, Sebastian Strempel, Andrew Hemphill, Norbert Müller, Junhua Wang, Bruno Gottstein, Markus Spiliotis

## Abstract

In this study, we present a comprehensive analysis of the hepatic miRNA transcriptome in mice suffering from experimental primary alveolar echinococcosis (AE), a parasitic infection caused upon ingestion of *Echinococcus multilocularis* (*E. multilocularis*) eggs. At one month post-infection, infected *C57BL/6* mice, along with non-infected control mice, were euthanized. Subsequently livers were collected and used for small RNA library preparation and next-generation sequencing (NGS). The most significantly dysregulated hepatic miRNAs were validated by Stem-loop RT-qPCR. We identified 28 miRNAs with significantly altered expression levels upon infection with *E. multilocularis*. Of these, 9 were up-regulated (fold change (FC) ≥ 1.5) and 19 were down-regulated (FC ≤ 0.66) as compared to the non-infected controls. In infected liver tissues, mmu-miR-148a-3p and mmu-miR-101b-3p were 8- and 6-fold down-regulated, respectively, and the expression of mmu-miR-22-3p was reduced by 50%, compared to non-infected liver tissue. Conversely, significantly higher hepatic levels were noted for *Mus musculus* (mmu)-miR-21a-5p (FC = 2.3) and mmu-miR-122-5p (FC = 1.8). Down-regulated miRNAs were highly enriched in Reactome and KEGG pathways of angiogenesis and fatty acids biosynthesis. Moreover, relative mRNA expression levels of three pro-angiogenic (VEGFA, MTOR and HIF1-*α*) and two lipogenic (FASN and ACSL1) genes were significantly higher in livers of *E. multilocularis* infected mice. Lastly, we studied the issue related to functionally mature arm selection preference (5p and/or 3p) from the miRNA precursor and showed that 9 pre-miRNAs exhibited different arm selection preferences in normal versus infected liver tissues. Our study provides first evidence of miRNA involvement in liver pathogenesis during AE. Our future research will focus on the characterization of miRNA transcriptome patterns in more advanced AE-stages towards the assessment of microRNA therapy for AE, and experimentally address functional characteristics of selected features presently found.

**Author Summary:** Various infectious diseases in humans have been associated with altered expression patterns of microRNAs (miRNAs), a class of small non-coding RNAs involved in negative regulation of gene expression. Herein, we revealed that significant alterations of miRNA expression occurred in murine liver subsequently to experimental infection with *Echinococcus multilocularis* (*E. multilocularis*) eggs when compared to non-infected controls. At the early stage of murine AE, hepatic miRNAs were mainly downregulated. Respective target genes of the most extensively downregulated miRNAs were involved in angiogenesis and fatty acid synthesis. Indeed, angiogenic and lipogenic genes were found to be significantly higher expressed in *E. multilocularis* infected livers relative to non-infected controls. These boosted cellular pathways are advantageous for development of the *E. multilocularis* metacestodes, since this larval stage is not able to undertake *de novo* fatty acid synthesis, and angiogenesis allows the larvae to be periparasitically supplied by oxygen and nutrients and to get rid of waste products. More research on the miRNA transcriptome at more advanced infection-stages, and on the role of angiogenesis in *E. multilocularis* larval growth and metastasis, is required to assess the usefulness of microRNA- and anti-angiogenic therapies against *E. multilocularis* infection.

## Introduction

Human alveolar echinococcosis (AE) is a parasitic disease caused by infection with the larval stage (metacestode) of the cyclophyllid tapeworm *Echinococcus multilocularis* (*E. multilocularis*; Cestoda, Taeniidae) [1]. In Europe, *E. multilocularis* undergoes a sylvatic life cycle that predominantly includes the red fox (*Vulpes vulpes*) as major definitive host (HD) and rodents (family Arvicolidae) acting as intermediate hosts (IHs) [2]. For humans, accidental infection with *E. multilocularis* eggs through the oral route can lead to the development of AE, affecting primarily the liver in 98% of cases [1]. Within the liver tissue, the asexual proliferation of metacestodes occurs by exogenous budding of new vesicles, thus the larval mass progrediently invades the surrounding hepatic tissue, with a potential of metastasis formation in distant sites such as lungs, brain and other organs [3–6]. Disease progression is largely supported by the ability of *E. multilocularis* metacestodes to modulate immunological host-defense mechanisms [7, 8].

AE is listed as one of the rare and neglected tropical diseases by the World Health Organization, and if left untreated the disease results in mortality in more than 90% of cases [9]. Radical surgical excision of the parasite tissue complemented by adjuvant chemotherapy is the only curative treatment for hepatic AE, but this applies to only ∼ 30% of patients [10]. For inoperable cases, the only currently licensed chemotherapeutical option is based on the benzimidazole derivatives albendazole and mebendazole. Benzimidazole-therapy has increased the survival rate of affected patients to 85-90% [4]. However, these drugs exhibit a parasitostatic rather than parasitocidal activity; hence patients must take these drugs lifelong. No viable alternatives to benzimidazole-based chemotherapy exist for patients suffering from benzimidazole intolerance [11]. The claim for a better management and control of AE calls for new treatment concepts, thus highlighting once more the necessity to gain deeper insights into the molecular basis of AE-induced liver pathology.

MicroRNAs (miRNAs) are a class of 21–24 nucleotides (nt) small non-coding RNAs discovered in the early 1990s as key regulatory factors of developmental timing in *Caenorhabditis elegans* [12]. The biogenesis of miRNAs is a two-step process; it begins in the nucleus where a miRNA gene is transcribed to primary miRNA (pri-miRNA) which will be processed by nuclear RNase III Drosha-like nucleases to generate a precursor hairpin miRNA (pre-miRNA). This pre-miRNA is then exported to the cytoplasm to become a mature miRNA [13, 14]. A miRNA can specifically bind to its target mRNA at the 3’ untranslated regions (3’UTRs), 5’ UTRs, exons and/or introns [15, 16]. This results in repression of the target mRNA expression by diverse mechanisms, including inhibition of the translational machinery, disruption of cap–poly (A) tail interactions, and exonuclease-mediated mRNA cleavage [17]. Thus, miRNAs are major elements of negative post-transcriptional regulation of gene expression. To date, more than 4000 miRNA-generating loci have been annotated in the human genome, representing thus 1 ─ 5% of all human genes [18–20], and more than 60% of human protein-coding genes harbor at least one conserved miRNA-binding site [21, 22]. Several cellular and biological processes such as cell proliferation, metabolism, apoptosis and immune defenses are orchestrated by miRNAs [23–25].

Alterations in microRNA gene expression have been reported for a wide range of human pathologies such as cancers, metabolic disorders and cardiovascular diseases [26, 27]. In hepatocellular carcinoma (HCC), dysregulated miRNAs have been assessed as drug targets [28, 29], disease stage / survival rate predictors [30] and as markers of responsiveness to therapy [31].

In recent years, significant efforts have been made in outlining and defining the roles of miRNAs in host-pathogen interactions, with a main focus on host miRNAs. In this context, evidence was provided that hepatitis C virus replication is completely dependent on the liver-specific miR-122 [32]. Similarly, changes of host-miRNA expression profiles have been associated to different helminthiases [33–36] where miRNAs dysregulation was relevant to tissue dysfunction and to the type of immune response [37]. In the case of *Schistosoma japonicum*, high serum level of hepatic miR-223 was correlated with active infection, and responsiveness to praziquantel therapy was characterized by a return to normal levels [38]. *E. multilocularis* was also found to quantitatively modulate circulating and liver miRNAs in a mouse model of secondary (intraperitoneal) infection [39, 40]. Similarly, the closely related *E. granulosus* also caused changes in the intestinal miRNA transcriptome of Kazakh sheep [41]. To date, miRNA-directed therapy against helminthiases is highly appealing [42]; both parasite - and host-derived miRNAs can be targeted, which allows (i) to interfere in essential biological processes of the parasite [43–45] and (ii) to ensure that the host environment returns to a normal biological state [46].

Growth of *E. multiocularis* larvae induces changes to liver metabolism that collectively result in a net mobilization of glucose, lipid and amino acids [47, 48]. Other studies demonstrated deviations in hepatic gene expression at early stage of experimental primary *E. multilocularis* infection in the murine model compared to non-infected liver tissue [49, 50]; however, how and whether miRNAs are involved in post-transcriptional regulation of gene expression remains unknown. In immunocompetent individuals, human AE usually takes decades before symptoms arise, and once diagnosed, patients often have already reached an advanced stage of disease hampering the prospect to achieve complete parasite clearance. Accordingly, understanding of early cellular and molecular events that take place along with intra-hepatic establishment of *E. multilocularis* larvae is a crucial task for the development of novel means of prevention and treatment.

In this study, we applied Illumina next-generation sequencing (NGS) to comprehensively analyze the miRNA expression profile in the mouse liver at the early stage (one month post-infection) of primary AE, and compared miRNA expression levels in infected livers to non-infected tissue samples. Results were validated based on quantitative stem-loop RT-PCR. Hepatic miRNAs that exhibited significantly altered disease specific expression levels were further studied by Reactome and KEGG enrichment analyses. Subsequently, for the top ranked cellular processes, we comparatively measured the relative mRNA levels of key associated genes in infected and non-infected livers tissue samples

This is the first survey of miRNAs regulation in early AE, which will contribute to a better understanding of the role of hepatic miRNAs in promoting parasite survival and growth.

## Materials and Methods

### Ethics statement

Mice were housed and handled under standard laboratory conditions and in agreement with the Swiss Animal Welfare Legislation (animal experimentation license BE 103/11 and BE 112/14).

### Mouse model and primary AE infection

Ten 8-week-old female *C57BL/6* mice were obtained from Charles River GmbH, Germany. The animals were divided into two groups, five animals each. The control group remained uninfected, and the other group was orally infected with *E. multilocularis* eggs. The eggs used in this study were obtained from a naturally infected fox that was shot during the official Swiss hunting season. To prepare the parasite eggs for subsequent infection of mice, the fox intestine was removed under appropriate safety precautions and cut into 4 pieces. After longitudinal opening of the intestinal segments, the worm-containing mucus was scraped out and put into petri dishes containing sterile water. Subsequently, the mucosal suspension was serially filtered through a 500µm and then 250µm metal sieve, by concurrently disrupting the worms with an inversed 2 ml syringe top. This suspension was further filtered through a 105 μm nylon sieve. The eggs were then washed by repeated sedimentation (1xg, 30 min., room temperature) in sterile water containing 1% Penicillin/Streptomycin and stored in the same solution at 4 °C. The sodium hypochlorite resistance test was used to assess egg viability [51], and to ascertain efficiency of infectivity, a preliminary test was done in two female *C57BL/6* mice. After confirmation of infectivity, five mice were each inoculated with approximately 1×10^3^ eggs suspended in 100 µl sterile water by gavage. The control mice received sterile water only. At one month post-infection, all animals were euthanized, and their livers were removed under sterile conditions for further analyses. All animal infections were performed in a biosafety level 3 unit (governmental permit no. A990006/3). One mouse from the uninfected control group died during the experiment from unknown causes and was thus not included in this experiment.

### Liver samples, total RNA extraction, small RNA library preparation and NGS

Liver tissue samples from *E. multilocularis-*infected mice were taken from the peri-parasitic area, precisely 3 to 4 mm adjacent to parasite lesions, which appeared as small white or yellowish spots. Liver tissue samples from uninfected animals were collected from the same hepatic areas. The obtained liver tissues were minced quickly, mixed at a ratio of 1:10 with QIAzol lysis reagent (Qiagen, Cat: 79306) and homogenized by bead beating (FastPrep-24 Instrument, MP Biomedicals, Cat: 116004500). Total RNA was extracted according to the manufacturer’s instruction of the QIAzol lysis reagent with the exception that chloroform was replaced by 1-bromo-3-chloropropane for the phase separation step (Sigma, cat: B9673). To remove genomic DNA contamination in RNA samples, an enzymatic digestion step using DNase I was carried out. Finally, total RNA was re-suspended in RNase-free water. RNA quantity and RNA quality number (RQN ≥ 8) was determined by the Fragment Analyzer CE12 (Advanced Analytics).

Two to five RNA extractions were prepared simultaneously from each liver, in order to subsequently choose the preparation with the best RQN. Samples from the infected mouse group were named as follows; AE-1pm-1.1, AE-1pm-2.1, AE-1pm-3.2, AE-1pm-4.1 and AE-1pm-5.1. Total RNA preparations from the uninfected control group were labelled ctr-1pm-1.1, ctr-1pm-3.2, ctr-1pm-4.2 and ctr-1pm-5.1 (first digit is the number of the mouse within its corresponding group and the second digit indicates the number of RNA preparation). Sequencing library construction and sequencing itself were performed by Microsynth (Balgach, Switzerland). Briefly, five libraries were generated from six mice (three mice from *E. multilocularis*-infected group and two mice from the uninfected control group). The total RNA was checked on an Agilent 2100 Bioanalyzer instrument for degradation. Subsequently, the CleanTag Ligation Kit (TriLink BioTechnologies) was used to prepare small RNA stranded libraries from total RNA (1µg RNA per library). Libraries were analyzed a second time on the Bioanalyzer to check for the intended miRNA fragment peak at 141 bp and subsequently a Sage Science Pippin Prep instrument was used to select the intended fragment size range. Prior to deep sequencing, the quality and concentration of each final library was assessed by PicoGreen measurements. The so refined libraries were then sequenced on an Illumina NextSeq 500 platform using a high-output v2 Kit (75 cycles) and targeting for an output of 50 Mio past-filter reads.

### Bioinformatics analysis of small RNA sequencing data

In a preprocessing step, reads were subjected to de-multiplexing and trimming of the TriLink adapter residuals using Illumina bcl2fastq v2 analysis software (bcl2fastq2 Conversion Software v2.19.1.). Quality of reads was checked with the software FastQC (v. 0.11.5) (https://www.bioinformatics.babraham.ac.uk/projects/fastqc/); thus, reads shorter than 10 bases or longer than 25 bases, or reads containing any “N” base, were discarded to refine the input for the statistical analysis. In a next step, clean reads of samples derived from the same experimental group (AE-1pm or ctr-1pm) were pooled together and then dereplicated using the software usearch (v. 8.1.1681) [52]. This resulted in a list of unique sequences that were annotated and recorded based on their frequency of occurrence. Within each dataset (AE-1pm and ctr-1pm), sequences having a read count of at least 10 were blasted against the miRBase [53, 54] mature miRNA mouse sequences (*Mus musculus*, mouse mm10 reference genome). Sequences that did not show a hit with an e-value of 1e-6 or smaller were subject to blast against the Rfam collection of RNA families (v. 12.3) [55]. In the latter search, hits with an e-value of less than one were regarded as valid for the sake of getting an overview. Mapping of cleaned reads to the mouse genome (v. mm10) was performed using the RNA mapping software STAR (v. 2.5.1b) [56]. Major parameters used were a minimum of 16 matching bases between read and reference, whereas a maximum of 1 mismatch was allowed and splice aware mapping switched off. Raw counting of uniquely mapped reads to annotated mature miRNAs was done with the software htseq-count from the HTSeq software suite (v. 0.6.1p1) [57]. The miRNA annotations stem from miRBase matching the mm10 reference genome. Normalization of the raw miRNA read counts and differential miRNAs expression analysis were conducted using the R package DESeq2 (v. 1.12.4); the DESeq2 method combines shrinkage estimation for dispersions and fold changes (log2FC) which allow a more sophisticated quantitative analysis [58, 59]. Two-dimensional principal component analysis (2D-PCA) [60] was applied to check whether control mice (ctr-1pm) could be distinguished based on their miRNA expression profiles from those experimentally infected with *E. multilocularis* eggs.

### Stem-loop reverse transcription (RT) and real-time quantitative (Stem-loop RT-qPCR) of dysregulated mature miRNAs

To validate the expression profile of the most dysregulated miRNAs obtained from the analysis of NGS data, we assessed the relative expression levels of twelve miRNA by stem-loop RT-qPCR, as previously described [61]. The small nucleolar RNA 234 gene (*sno234*) was included as reference for microRNA normalization [62]. In total, 9 liver tissue samples derived from the AE-infected (5 mice) and uninfected mice (4 mice) were individually assessed.

Briefly, 1 μg of total RNA template was reverse transcribed to cDNA by M-MLV reverse transcriptase (Promega) using specific stem-loop RT primer and under the following conditions: 5 min_25°C, 60 min_42°C, 15 minutes_70C, reactions were then cooled down to 4°C. All q-PCRs were performed in Rotor-Gene 6000 (Corbett Life Science) and using FastStart Essential DNA Green Master Kit (Roche, Switzerland). The total volume for qPCR was 10 μl, consisting of 5 μl of Faststart Essential DNA Green Master Mix (2xconc), 1 μl of forward and reverse primers, and 2.5 µl of cDNA template (diluted 1:5). All experiments were run in triplicates. Quantitative PCRs were performed according to the following program: an initial hold at 95°C for 15 min, 40 cycles (94°C for 15 s, annealing at 63°C for 20 s, extension at 72°C for 20 s and a final denaturation step from 50 °C to 95 °C. Specificity of each qPCR and presence of primer dimers were checked based on analysis of the generated melting curve. All primers characteristics are listed in Table 1.

**Table 1:**
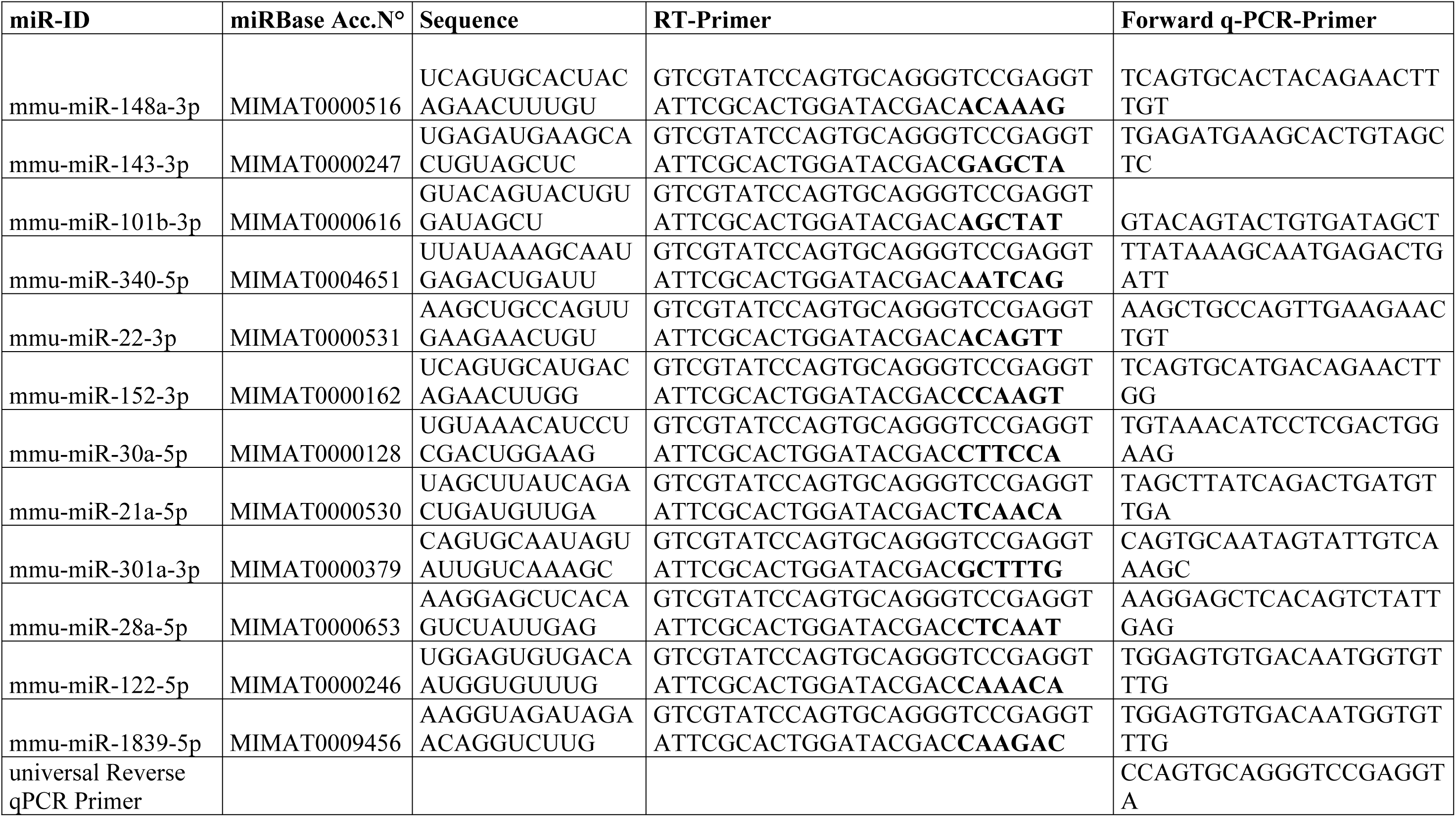
Stem-loop RT-qPCR primers

For relative miRNA expression, data were expressed as median ± standard deviation (SD) and examined for statistical significance with the nonparametric Mann–Whitney U test. *P*-values of less than 0.05 were considered to be statistically significant.

### MicroRNA target prediction and pathway enrichment analysis

Significantly differentially expressed miRNAs with FC ≥ 1.5 or FC ≤ 0.66 were chosen for target prediction. Since miRNAs inhibit their target mRNAs, reduction of miRNA expression lead to up-regulation of the target genes and vice versa. For miRNA-mRNA target predictions, we used miRNet [63], a database for network-based visual analysis of miRNAs, targets and functions. MiRNet integrates high-quality miRNA-target interaction databases (miRTarBase v6.0, TarBase v6.0 and miRecords); these databases provide direct experimental evidence regarding the miRNA–target interaction. For functional and pathway enrichment analysis, we used two pathway databases, including Reactome [64] and KEGG [65]. Statistical significance was measured by applying hypergeometric test [63].

### Expression analysis of miRNA target genes using RT-qPCR

Since angiogenesis and fatty acids biosynthesis were the most enriched pathways of the downregulated miRNAs, we selected five key genes relevant to these identified pathways for further analysis of their mRNA levels in AE-infected- and uninfected mice (Table 2). Precisely, RT-qPCR was employed to assess the relative expression of three main pro-angiogenic factors, namely (i) vascular endothelial growth factor A (VEGFA); (ii) the mechanistic target of rapamycin (MTOR); and (iii) hypoxia inducible factor 1, α subunit (HIF1α). In addition, the expression of two lipogenic genes was examined, namely fatty acid synthase (FASN) and acyl-CoA synthetase long-chain family member 1 (ACSL1). All genes were assessed individually in five liver tissue samples from AE-infected and four samples from uninfected mice (same experiment). We used cDNA samples prepared for the Stem-loop RT-qPCR for validating the set of 28 dysregulated mature miRNAs (see section above). The qPCRs were performed as described above, with the exception that the annealing temperature was set at 62° C for all five genes. We used the *glyceraldehyde-3-phosphate dehydrogenase* (*gapdh*) as endogenous reference. Detailed information on qPCR primers are provided in Table 2.

**Table 2:**
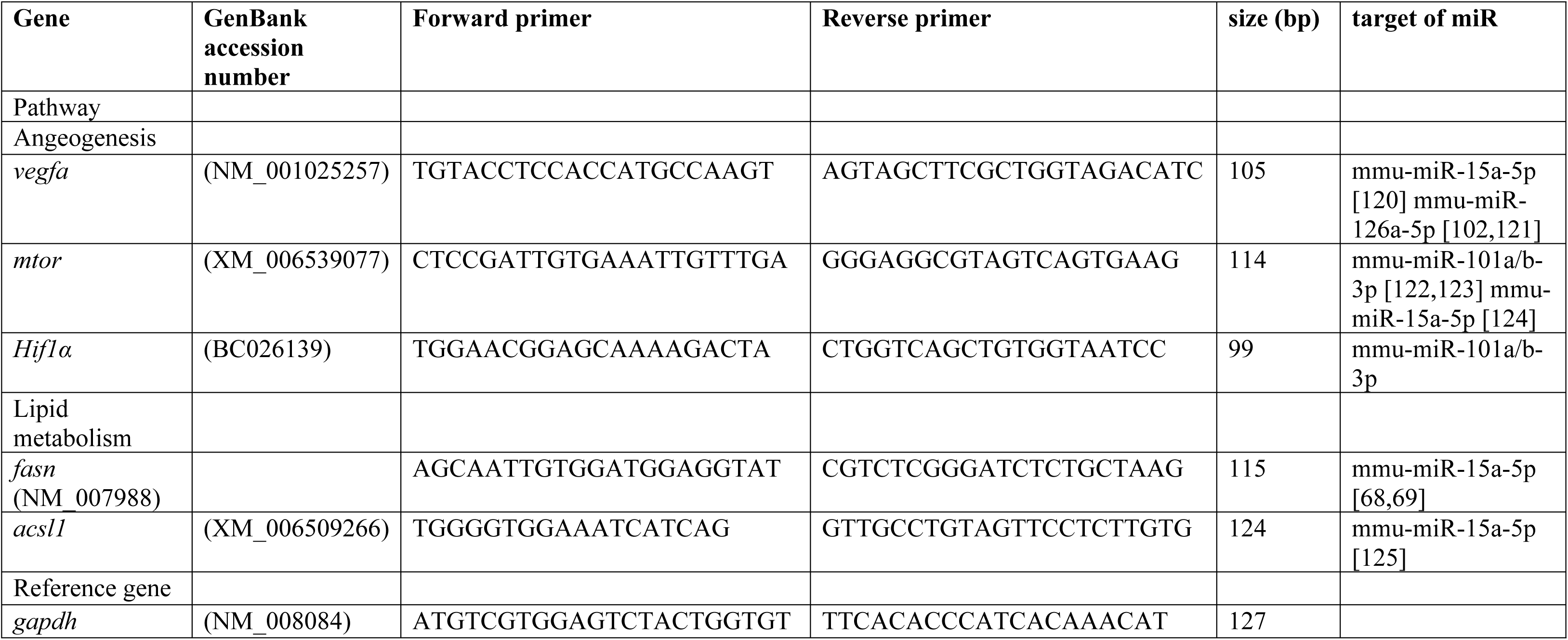
qPCR primers

## Results

### Next generation sequencing (NGS) data

To characterize the miRNA transcriptome in murine liver during early stage of primary AE, small RNA libraries from three *E. multilocularis*-infected and two uninfected control mice were constructed and subjected to high-throughput sequencing.

The number of uniquely mapped reads with an average mapped length of 21 nucleotides ranged from 4.831.114 to 1.749.597, representing thus 66 to 60% of the total cleaned reads. More than 95% of those uniquely mapped reads were assigned to miRBase annotated miRNAs (Table 3).

**Table 3:** Summary of raw counts of reads mapped to miRNA features of the reference sequence.

| Library | <sup>a</sup>Cleaned Reads | Uniquely Mapped Reads | Reads On Feature | Reads No Feature | Reads Ambiguous |
| --- | --- | --- | --- | --- | --- |
| AE-1pm-2.1 | 6802673 | 3840087 | 3659366 | 180718 | 3 |
|  |  |  | 95.3% | 4.7% | 0% |
| AE-1pm-3.2 | 7315799 | 4831114 | 4698825 | 132287 | 2 |
|  |  |  | 97.3% | 2.7% | 0% |
| AE-1pm-5.1 | 7098967 | 4591014 | 4459799 | 131210 | 5 |
|  |  |  | 97.1% | 2.9% | 0% |
| ctr-1pm-3.2 | 2901675 | 1749597 | 1690171 | 59426 | 0 |
|  |  |  | 96.6% | 3.4% | 0% |
| ctr-1pm-4.2 | 6683305 | 3861026 | 3704668 | 156358 | 0 |
|  |  |  | 96% | 4% | 0% |
| **<sup>a</sup>: reads considered for mapping** |  |  |  |  |  |

For all libraries, length distribution analyses revealed that length of most abundant sequences ranged from 18 to 24 nt with a peak at 21 nt (Fig 1A). In both groups, most miRNAs were detected with read counts below 100 (Fig 1B).

**Fig 1.**
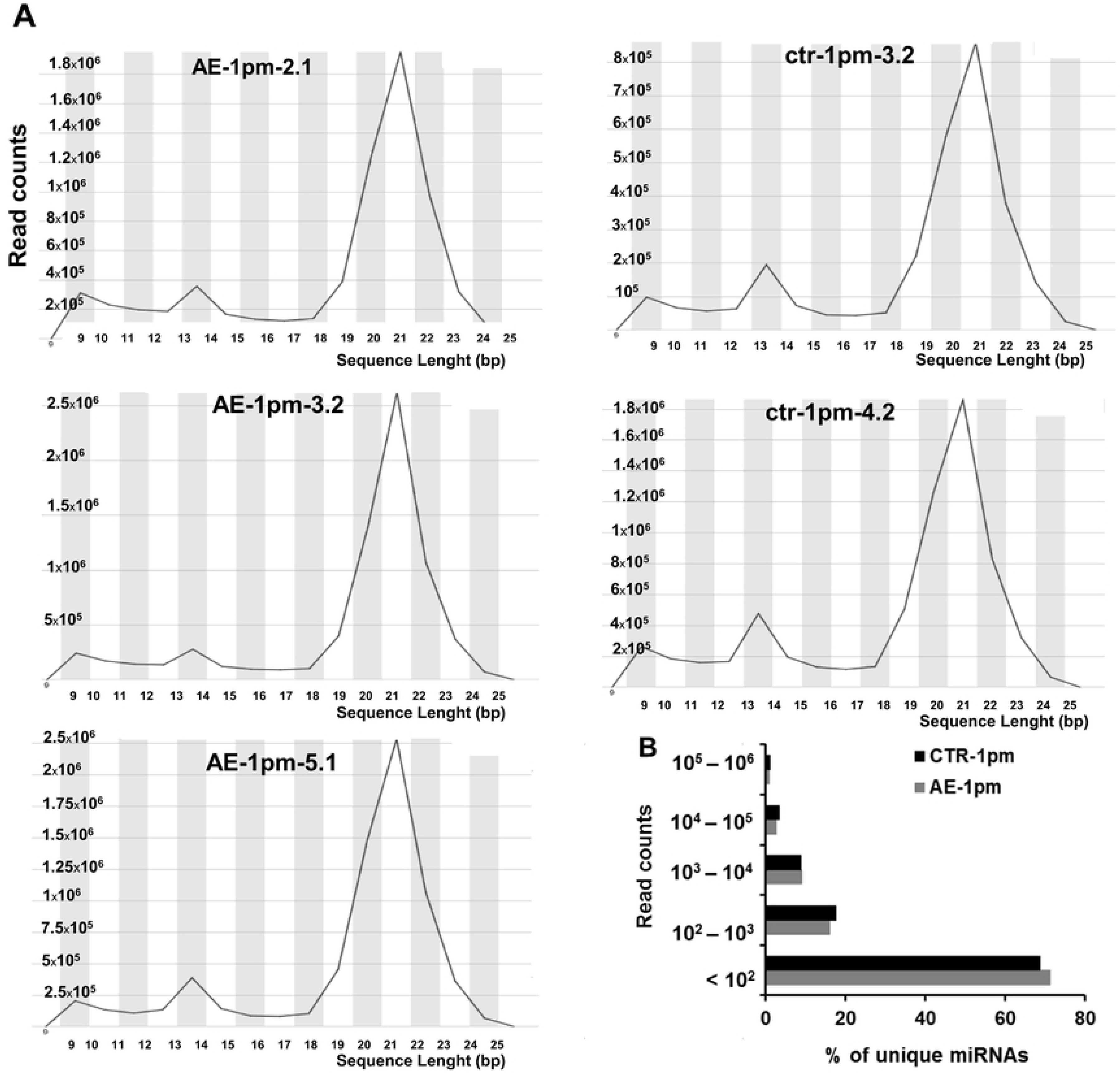
Characterization of the miRNAs identified by NGS from AE-infected and control animals. **(A)** Length distributions of total reads in the five small-RNA libraries. On the X-axis, reads < 10 nucleotides and reads > 25 nucleotides were discarded. The Y-axis depicts the read counts. The peak for the miRNA candidates (21 nucleotides) is centered. **(B)** The frequency of miRNAs that are expressed at the defined levels in each group, most miRNAs are expressed with a read count less than 100.

### Overview of miRNA expression profile in infected and uninfected mouse liver

A total of 699 known miRNAs were identified from both infected (AE-1pm) and uninfected (ctr-1pm) groups. Among these molecules, 530 were common to both groups with 124 and 45 miRNAs specifically expressed in the AE-1pm and ctr-1pm libraries, respectively. From the co-expressed miRNA cluster, a set of 87 miRNAs with a read count > 1000 in at least one of the two experimental groups was identified (Fig 2A). In liver-tissue samples derived from AE-infected mice, mmu-miR-122-5p, mmu-miR-21a-5p and mmu-miR-192-5p accounted for up than 70% of the total normalized miRNA counts, and they still compromised more than 50% total miRNA abundance in the control group (Fig 2B).

**Fig 2.**
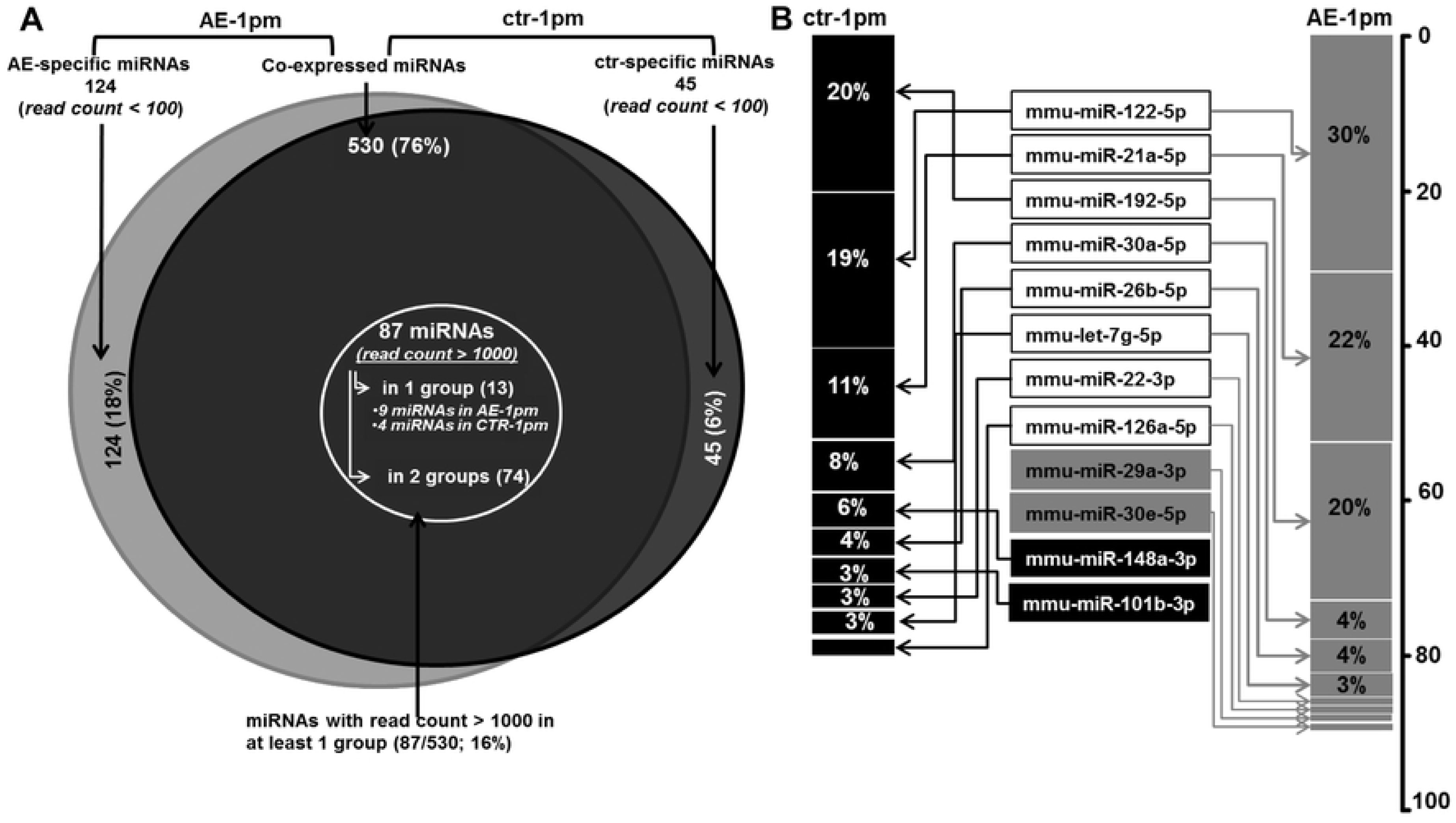
miRNA expression profile in livers of AE-infected and uninfected mice. **(A)** Venn diagram showing the overlap of expressed miRNA in liver-tissue samples from AE-infected and uninfected animals. **(B)** Top 10 most abundant miRNA and their frequency (%) in both groups.

### Alteration of the hepatic microRNA expression profile at early stage of hepatic AE

The 2D-PCA (Fig 3A) as well as the heatmap of sample-to-sample distances (Fig 3B) showed a clear separation of miRNA expression pattern between *E. multilocularis* egg infection versus healthy control. Globally, the observed clustering of samples from the same group indicated that a change in miRNAs expression pattern had occurred following infection.

**Fig 3.**
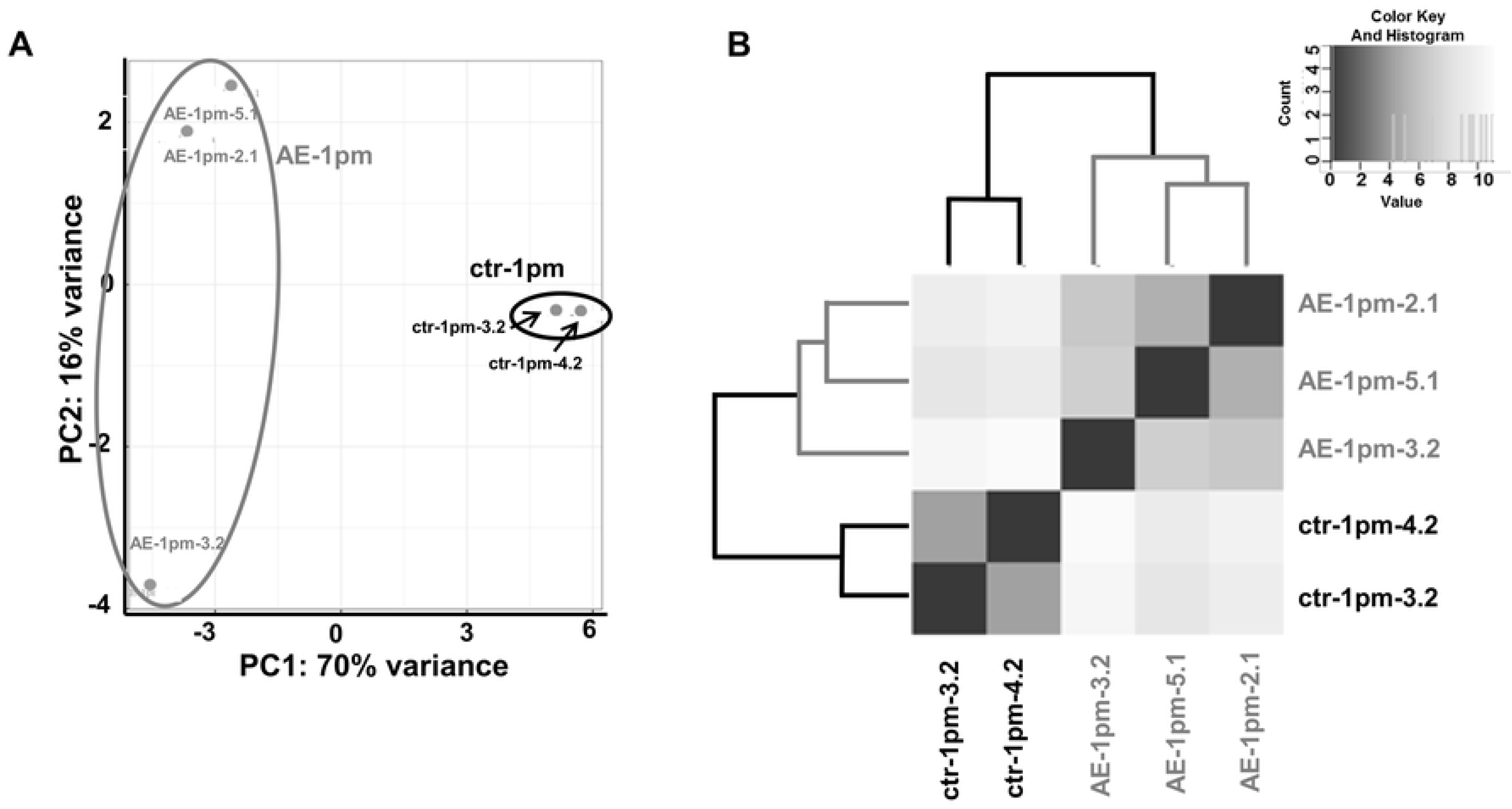
Principal component analysis (**A**) and (**B**) hierarchical clustering both revealed a clear separation between hepatic miRNA profiles from infected and uninfected mice.

A global comparative analysis of all miRNA read counts between both experimental mouse groups was carried-out and revealed the presence of a set of miRNAs whose log2FC was significant. From this latter cluster, we considered as significantly dysregulated miRNAs only those with (i) a normalized read count ≥ 1000, and (ii) FC ≥ 1.5 (Log2FC ≥ 0.58) or FC ≤ 0.6 (Log2FC ≤ −0.58). Thus, a total of 28 miRNAs were found to be differentially expressed in diseased livers compared to healthy controls (Fig 4). More information on detailed counts is shown in Table 4. The highest up-regulated miRNA in *E. multilocularis* infected livers was mmu-miR-21a-5p with a FC = 2.3. Conversely, the expression of mmu-miR-148a-3p was ∼8-fold lower as compared to control liver samples.

**Fig 4.**
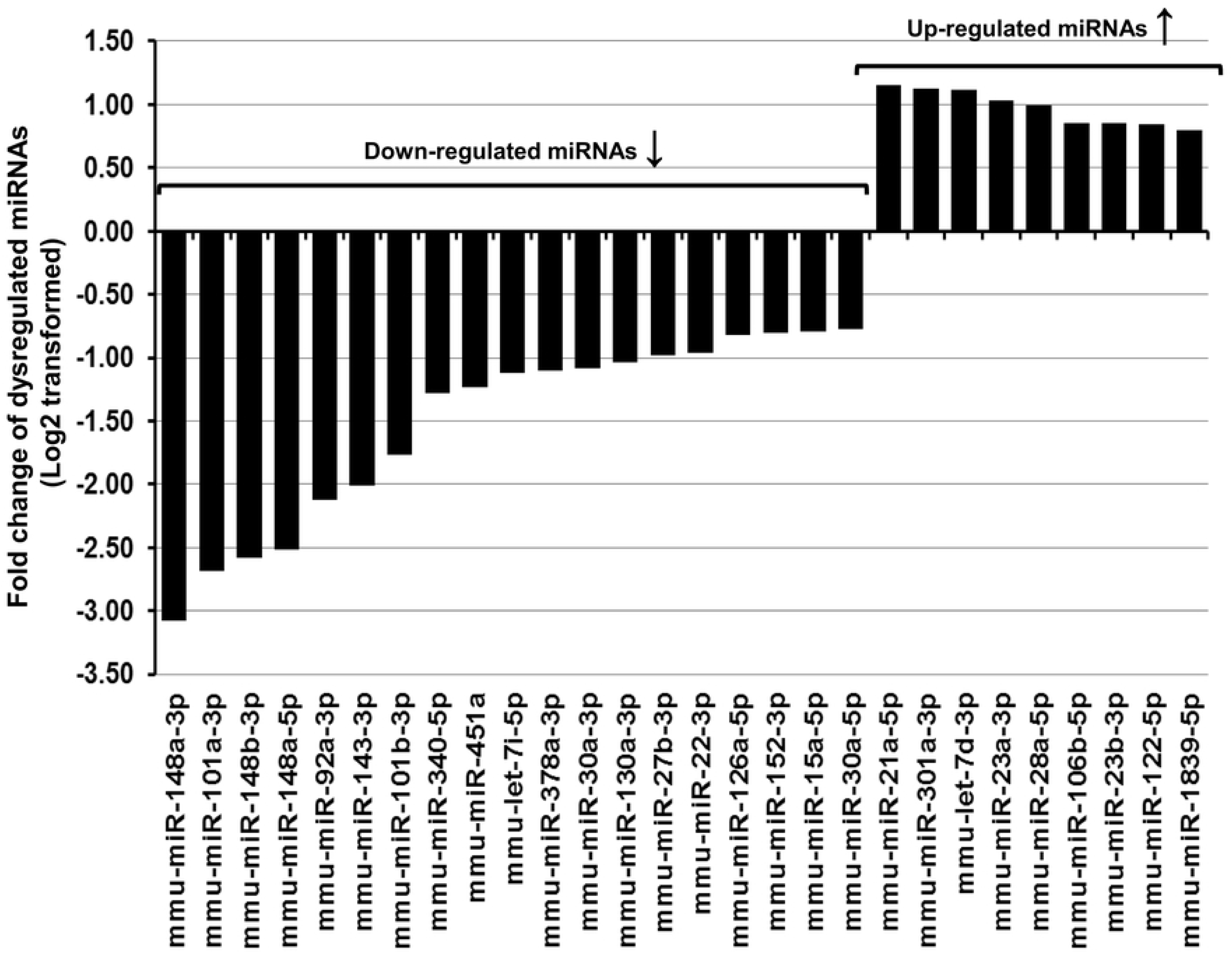
Twenty-eight dysregulated hepatic miRNAs in primary AE-infected mice compared to control healthy individuals: nine were up-regulated (Log2FC ≥ 0.58) and nineteen were down-regulated (Log2FC ≤ −0.58).

**Table 4:**
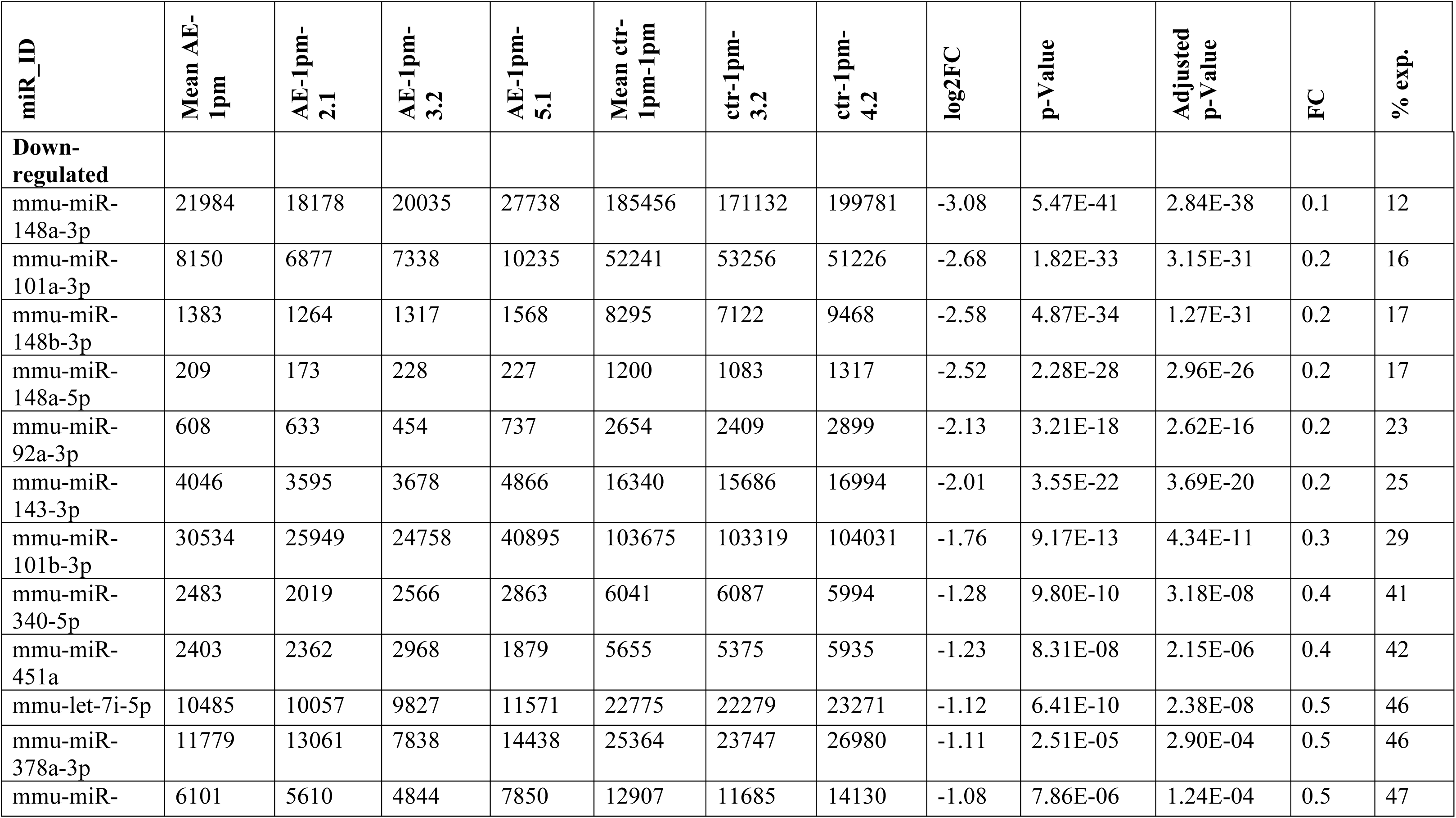
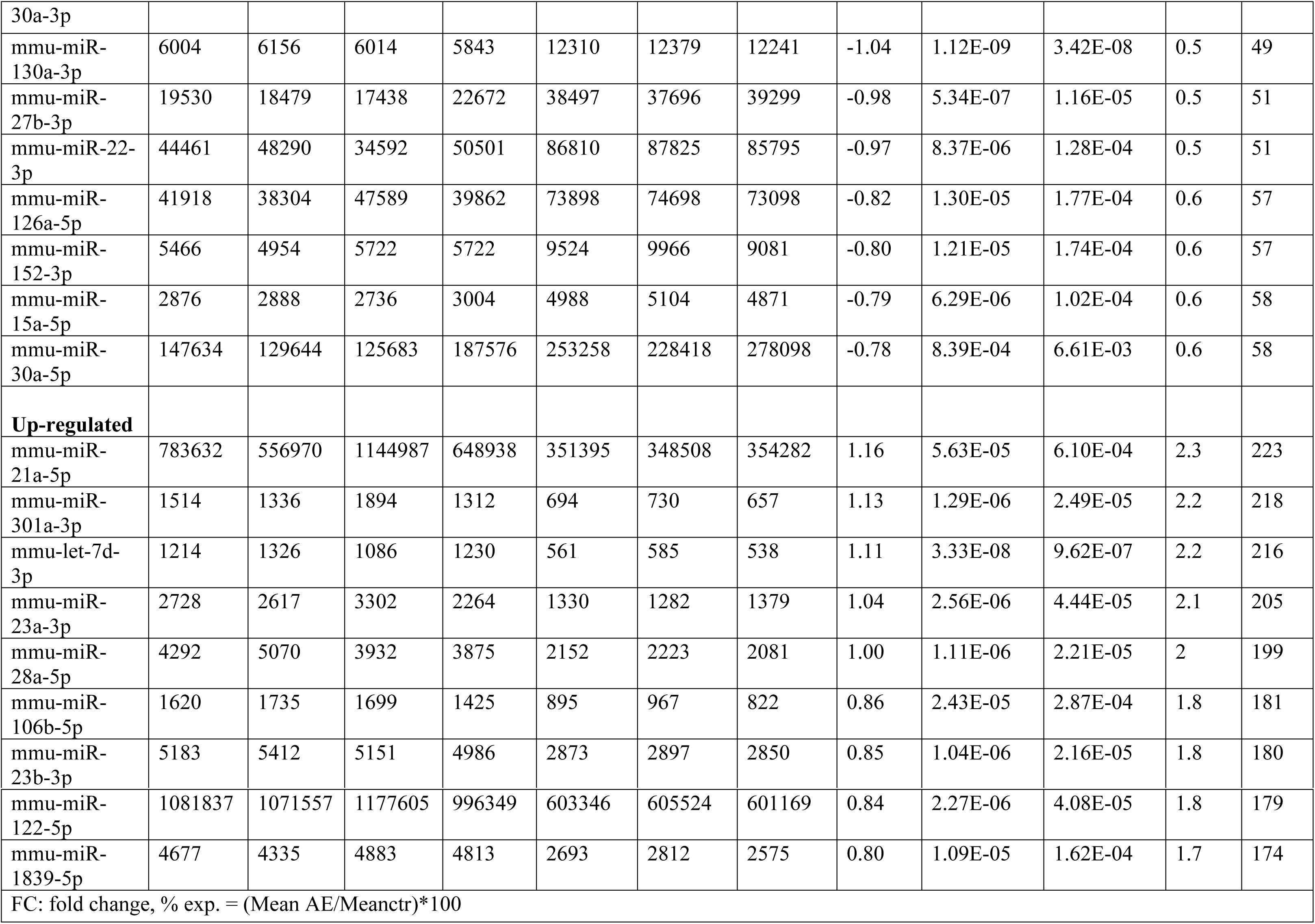
Dysregulated miRNAs in primary AE.

Stem-loop RT-qPCR was applied to validate the miRNA NGS data of 12 out of the 28 differentially expressed miRNAs. In order to optimize the statistical significance of this study, two additional samples were included into each group, thus the AE-1pm group comprised five animals and the ctr-1pm included four samples. Overall, the Stem-loop RT-qPCR results largely confirmed the results obtained through NGS as shown in Figure 5 (Fig 5). Seven miRNAs; mmu-miR-148a-3p, mmu-miR-143-3p, mmu-miR-101b-3p, mmu-miR-340-5p, mmu-miR-22-3p, mmu-miR-152-3p, mmu-miR-30a-5p were significantly less expressed in AE-1pm samples compared to ctr-1pm samples. In contrast, infected mice exhibited significantly higher expression levels of mmu-miR-21a-5p, mmu-miR-28a-5p, mmu-miR-122-5p and mmu-miR-1839-5p compared to the control mice. No significant difference in expression levels in the two groups were noted for mmu-miR-301a-3p.

**Fig 5.**
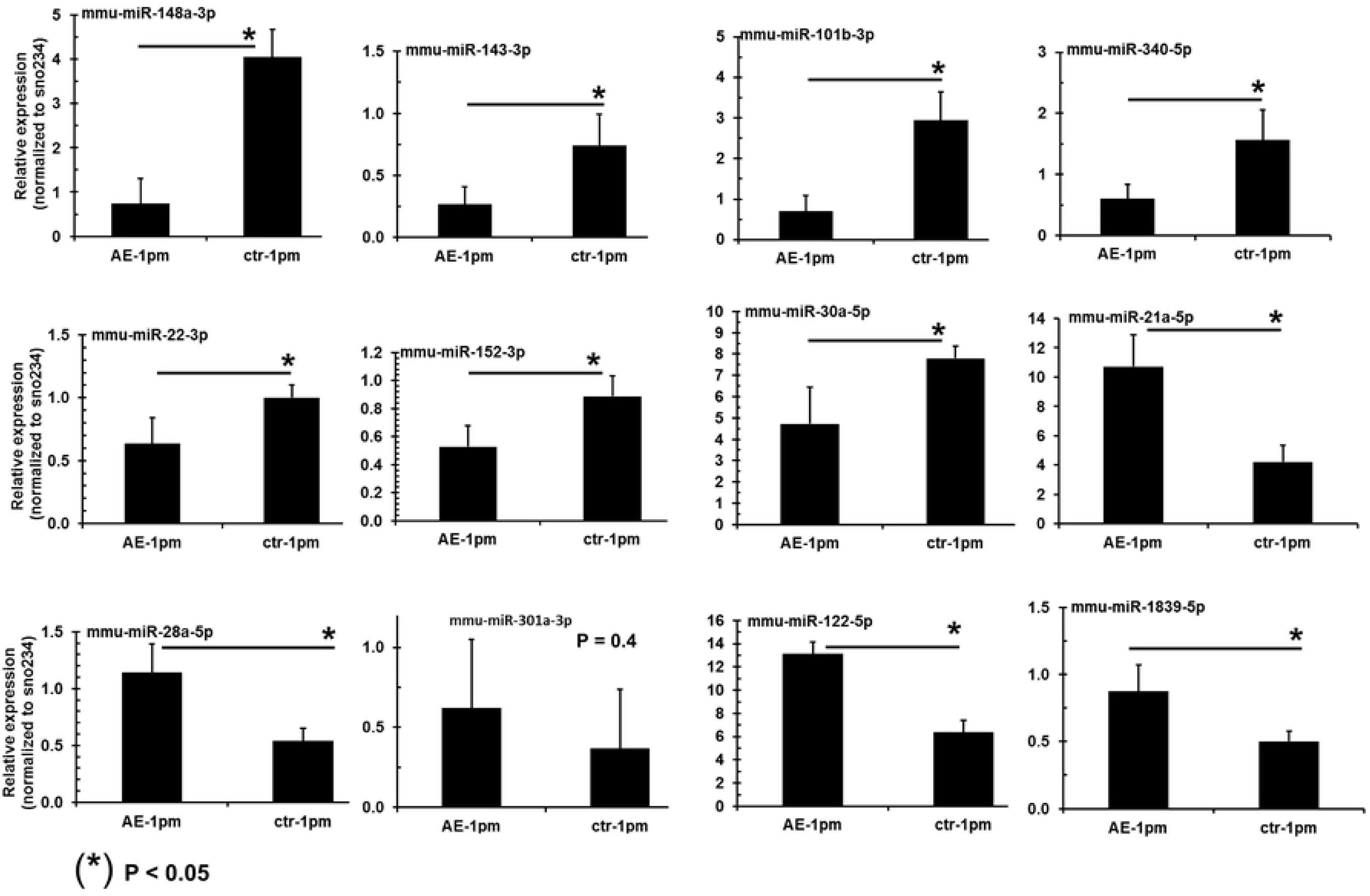
miRNAs expression validation by stem-loop RT-qPCR. Relative miRNA expression levels in livers of AE-infected and uninfected control mice. Normalization was done using *sno234* as the endogenous control. Bars represent median ± standard deviation. All stem-loop RT-qPCRs for each miR used biological replicates (n = 5 infected group and n = 4 uninfected group), with three technical replicates per experiment.

### miRNA precursors: preference selection of 5p- and 3p arm in *E. multilocularis*-infected– and uninfected liver tissues

In order to interrogate arm selection preferences of miRNA pairs (miRNA-5p/3p) in normal and infected liver tissues, we investigated the expression levels of miR-5p and miR-3p strands of the same miRNA precursor (pre-miRNA). We identified a total of 127 miRNA pairs that were co-expressed in all tissue samples and they exhibited a normalized read count ≥ 100 for 5p or 3p strand under infected and non-infected conditions. For each miRNA duplex, we calculated the selection rate (S) of 5p and 3p arm from the total read count (5p and 3p). For 25 (20 %) and 21 (17 %) out of these 127 miRNA pairs, mature miRNA was strictly derived from 3p (S%_miR-5p = 0; S%_miR-3p = 100) and 5p (S%_miR-5p = 100; S%_miR-3p = 0), respectively, and this was independent from the primary AE infection. Regarding the remaining 81 pre-miRNAs out of the total 127 identified pairs, mature miRNAs were derived from both arms. For 16 pre-miRNAs (out of the 81), 5p and 3p selection rates remained unchanged between infected and non-infected liver tissue, whereas selection preference for either the 5p- or the 3p-strand differed for the lasting 65 miRNA pairs (out of the 81).

Particularly for nine pre-miRNA (miR-106b, miR-144, miR-16-1, miR-1981, miR-214, miR-28a, miR-335, miR-345 and miR-532) the difference in 5p- and 3p-arm expression between the two groups ranged between 10% and 33% with *P-*value < 0.01 (Fig 6B). Overall, linear regression analysis of 5p-arm selection rate in the 127 miRNA pairs showed a strong correlation (R2 = 0.9764) between both groups (Fig 6A).

**Fig 6.**
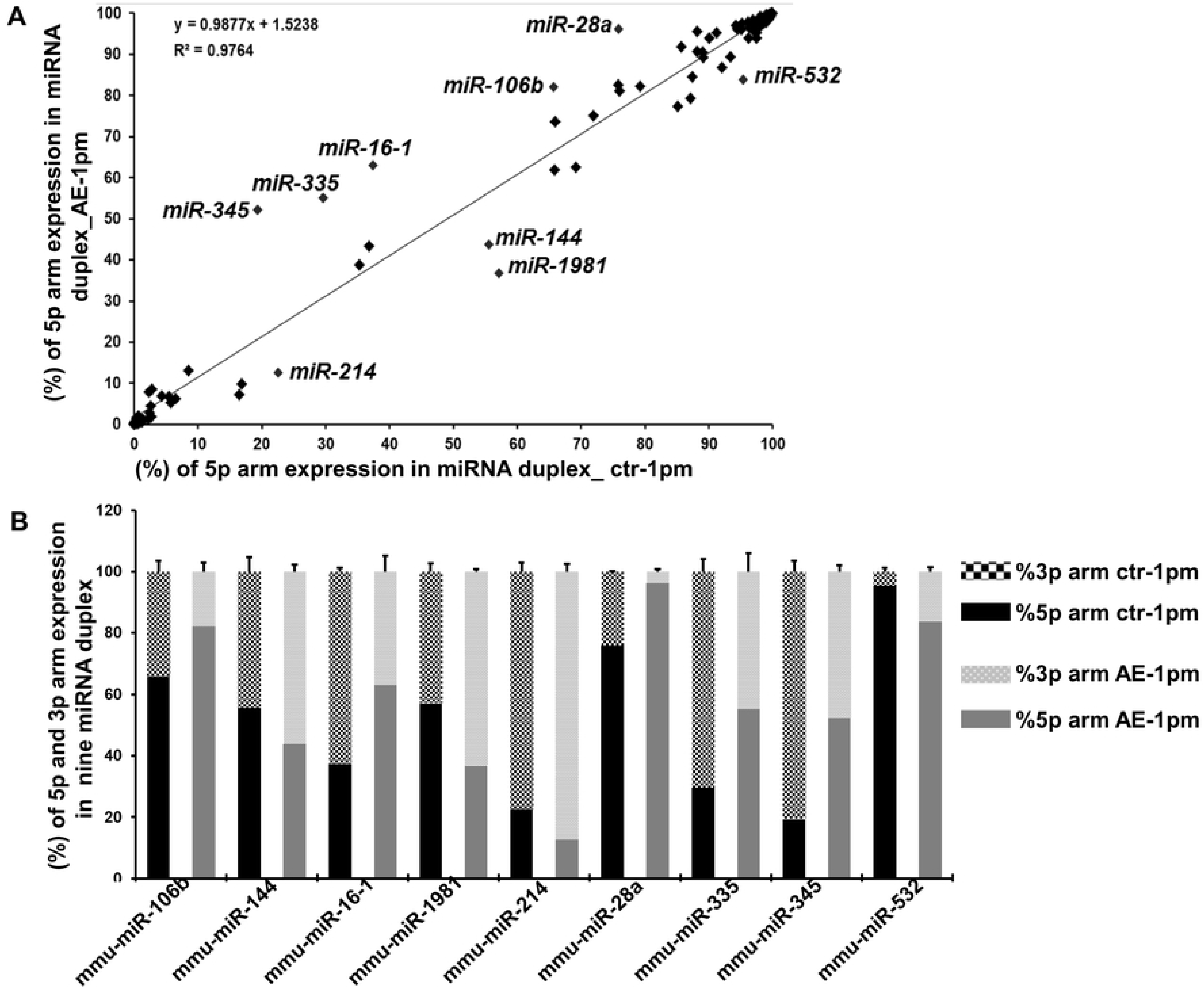
Arm selection preference (5p and 3p) in hepatic pre-miRNAs: AE-infection versus no-infection (control). (**A**) Linear regression analysis of the correlation in 5p-arm selection rate between *E. multilocularis*-infected (AE-1pm) and uninfected mice (ctr-1pm). The 5p-arm selection rate was determined for 127 miRNA pairs. (**B**) Nine pre-miRNAs exhibited a significant difference in 5p- and 3p-arm expression between *E. multilocularis*-infected (AE-1pm) and uninfected animals (ctr-1pm). For each of the nine miRNAs, the 100% stacked bars compare the 5p- and 3p-arm expression rate between AE-1pm and ctr-1pm mouse group.

### Prediction of the target genes of dysregulated host miRNAs

Potential targets of the 28 differentially expressed host miRNAs were predicted using the miRNet tool which was based on miRTarBase v6.0, TarBase v6.0 and miRecords algorithms. In total 1645 target genes were identified for the 25 miRNAs, while for the remaining three miRNAs (mmu-miR-148a-5p, mmu-miR-30a-3p and mmu-let-7d-3p), no target genes were found. Overall, from the 1645 targets, two clusters of unequal sizes were defined: a major group (1484/1645; 90%) includes target genes that are unique to one miRNA and a smaller set of 161 genes that are commonly controlled by two or more of the dysregulated miRNAs.

For the 17 down-regulated miRNAs, a set of 1426 target genes were found; almost 80% (1140/1426) of these miRNA-target interactions (MITs) were supported by experimental evidence. Among the 1426 target genes, 134 (10%) were shared by at least two miRNA molecules; out of these 134 targets, 49% (66/134) were common between mmu-miR-15a-5p and mmu-miR-340-5p.

On the other hand, for the eight up-regulated miRNAs, 219 target genes were identified, and a smaller number of MITs (117/219; 50%) were experimentally validated. Among the 219 identified target genes, 89 were regulated only by mmu-miR-122-5p, with 71 MITs being experimentally validated. In total, twenty six genes were common between two microRNAs; alone mmu-miR-23a-3p and mmu-miR-23b-3p share 21 targets, 81% of common genes.

The 1645 putative target genes covered a wide range of biological functions, notably those related to immunity, metabolism and epigenetic modifications such as DNA methylation and histone modification. Immune relevant genes included IL-1β targeted by mmu-miR-122-5p, SMAD3/4 transcription factors targeted by mmu-miR-27b-3p and mmu-miR-122-5p, calcium/calmodulin-dependent protein kinase II alpha (Camk2a) targeted by mmu-miR-340-5p, mmu-miR-148a-3p, mmu-miR-148b-3p and mmu-miR-152-3p, interferon regulatory factor (IRF) 7/8 targeted respectively by mmu-miR-122-5p and mmu-miR-22-3p, IL-17 receptor A (IL-17RA) targeted by mmu-miR-23a/b-3p, inducible T-cell costimulator (ICOS) targeted by mmu-miR-101a-3p and vascular cell adhesion molecule (V-CAM)-1 target by mmu-miR-340-5p. Two genes were directly involved in fatty acids (FAs) biosynthesis; acyl-CoA synthetase long-chain family member 1 (ACSL1) and Fatty Acid Synthase (Fasn) targeted by mmu-miR-340-5p and mmu-miR-15a-5p.

Relevant genes involved in epigenetic modifications included histone deacetylase (HDAC)-2/4/9 targeted by mmu-miR-340-5p, mmu-miR-22-3p and mmu-miR-340-5p, respectively, and DNA methyltransferases (Dnmt)-1, targeted by mmu-miR-148a/b-3p and mmu-miR-152-3p, respectively.

Interaction networks of miRNAs-targets were constructed (Supplementary Fig 1 and 2).

### Pathway enrichment analysis for dysregulated miRNAs

To get an overview on cellular pathways in which dysregulated miRNAs could be involved, the putative target genes were subjected to Reactome and KEGG for functional enrichment and pathway analysis. Analyses for down- and up-regulated miRNAs were made separately.

For down-regulated miRNAs: 56-Reactome- and 68 KEGG pathways (*P*-values < 0.01) were associated with up-regulated target genes. The vascular endothelial growth factor A-vascular endothelial growth factor receptor 2 (VEGFA-VEGFR2) and axon guidance were the most enriched pathways with up-regulated miRNA target genes on the basis of their statistical significance. Top 30 Reactome- and KEGG- -enriched pathways among up-regulated genes are shown in Figure 7 (Fig 7A and B). Among the 1426 up-regulated genes, 25 were involved in the Reactome pathway “VEGFA-VEGFR2” (Supplementary Fig 1).

**Fig 7.**
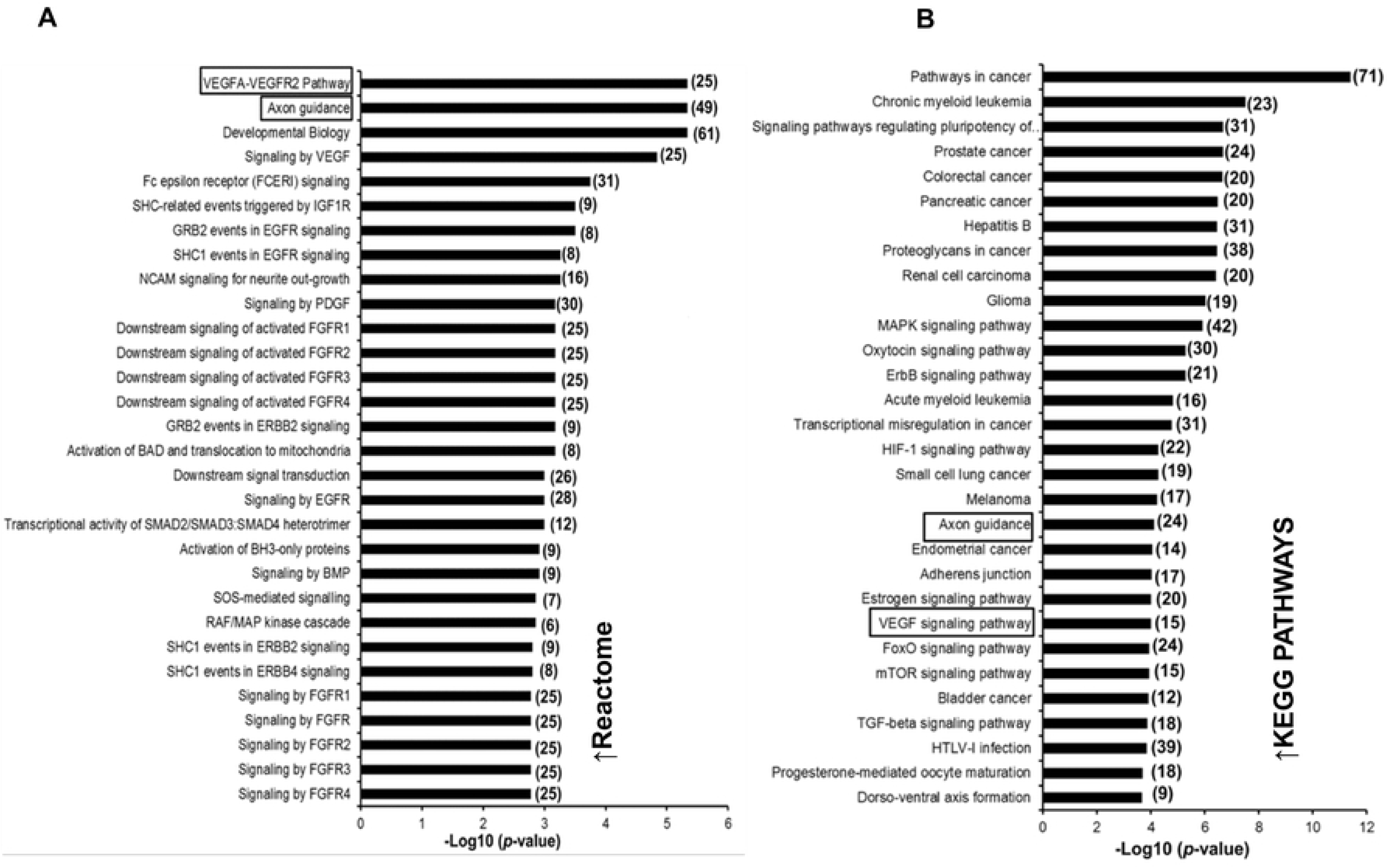
Functional analysis of predicted targets of down-regulated miRNAs. A and B: Top 30 overrepresented canonical pathways for up-regulated gene targets of down-regulated miRNAs according to Reactome and KEGG database, respectively. Pathways are ranked by score (-log10 (*P*-value). A higher score indicates that the pathway is more significantly associated with genes of interest. The numbers between brackets indicate the number of genes involved in each pathway.

Up-regulated miRNAs: 6-Reactome pathways and 23 KEGG pathways processes (*P*-values < 0.01) were associated with the down-regulated genes. As shown in Figure 8 (Fig 8 A and B), heme biosynthesis was the most affected pathway with seven involved genes, which are highlighted in Supplementary Figure 2.

**Fig. 8.**
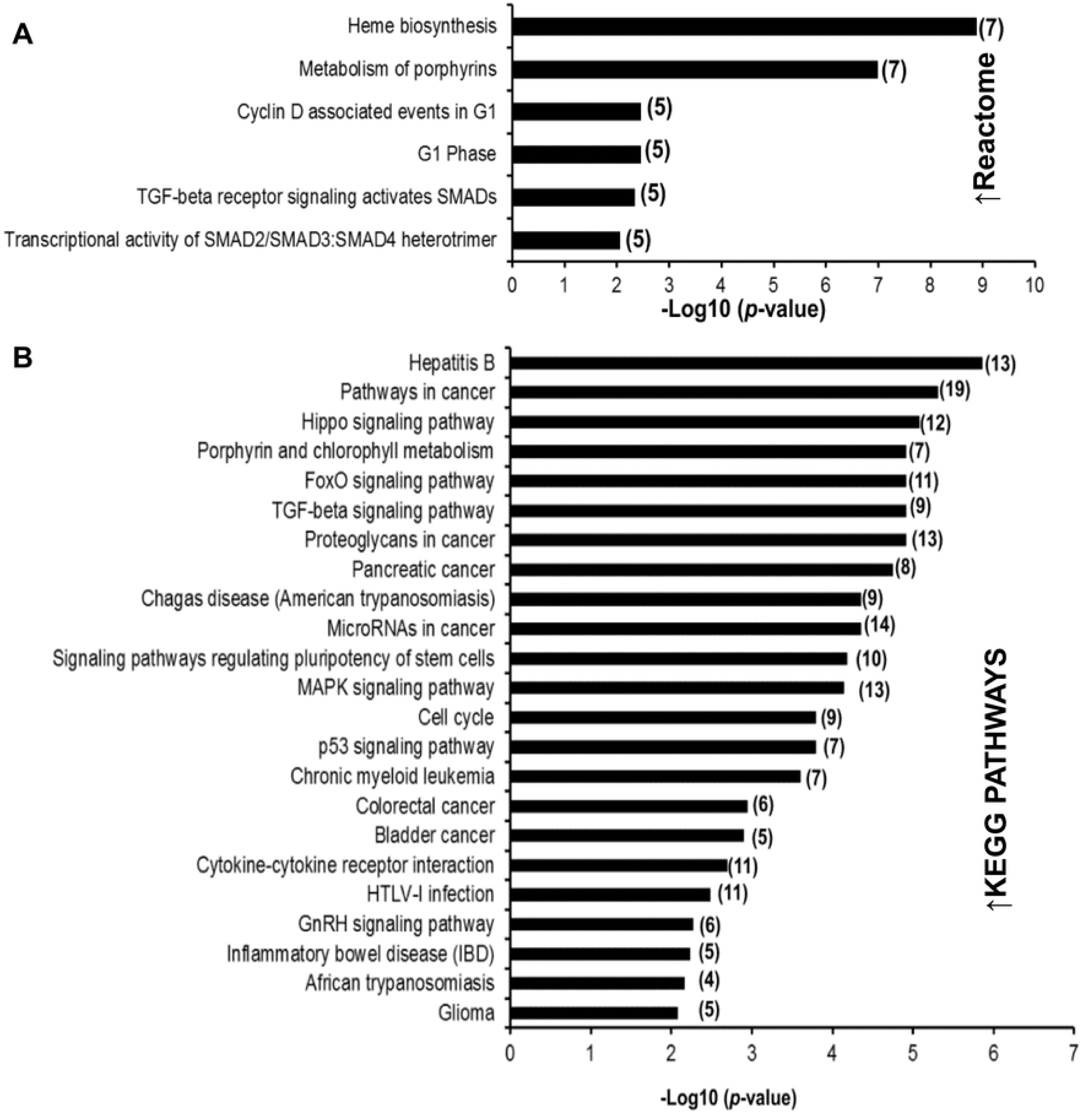
Functional analysis of predicted targets of down- and up-regulated miRNAs. A and B: Significant over-represented canonical pathways for down-regulated gene targets of up-regulated miRNAs according to Reactome and KEGG database, respectively. Pathways are ranked by score (-log10 (*P*-value). A higher score indicates that the pathway is more significantly associated with genes of interest. The numbers between brackets indicate the number of genes involved in each pathway.

### Expression analysis of key pro-angiogenic and fatty acid synthesis –associated genes as targets of down-regulated miRNAs

Since angiogenesis and fatty acid synthesis were the most significant cellular pathways that were enriched with target genes of down-regulated microRNAs, the relative expression of key genes involved in these cellular processes was further assessed. Thus five mRNA targets of down-regulated miRNAs were chosen using the following criteria: (1) predicted as a target of at least one of the down-regulated miRNA; (2) experimentally validated target of at least one of the down-regulated miRNA; (3) significantly relevant gene to the considered pathway; (4) a combination of 1, 2 and 3 (Table 2). Thus, the expression leves of three pro-angiogenetic (VEGFA, MTOR and HIF1α) and of two lipogenic (FASN and ACSL1) genes were comparatively assessed by RT-qPCR in livers from AE-infected and non-infected mice.

As shown in Figure 9, significantly higher mRNA levels for all 9 genes were detected in livers from *E. multilocularis*-infected mice relative to non-infected control group (Figure 9), which in turn confirmed that decreased miRNA expression was associated with increased target mRNA expression levels. Besides, these results strongly suggested that *E. multilocularis* egg infection could lead to increased angiogenesis and modulated fatty acid biosynthesis in the murine liver.

**Fig. 9.**
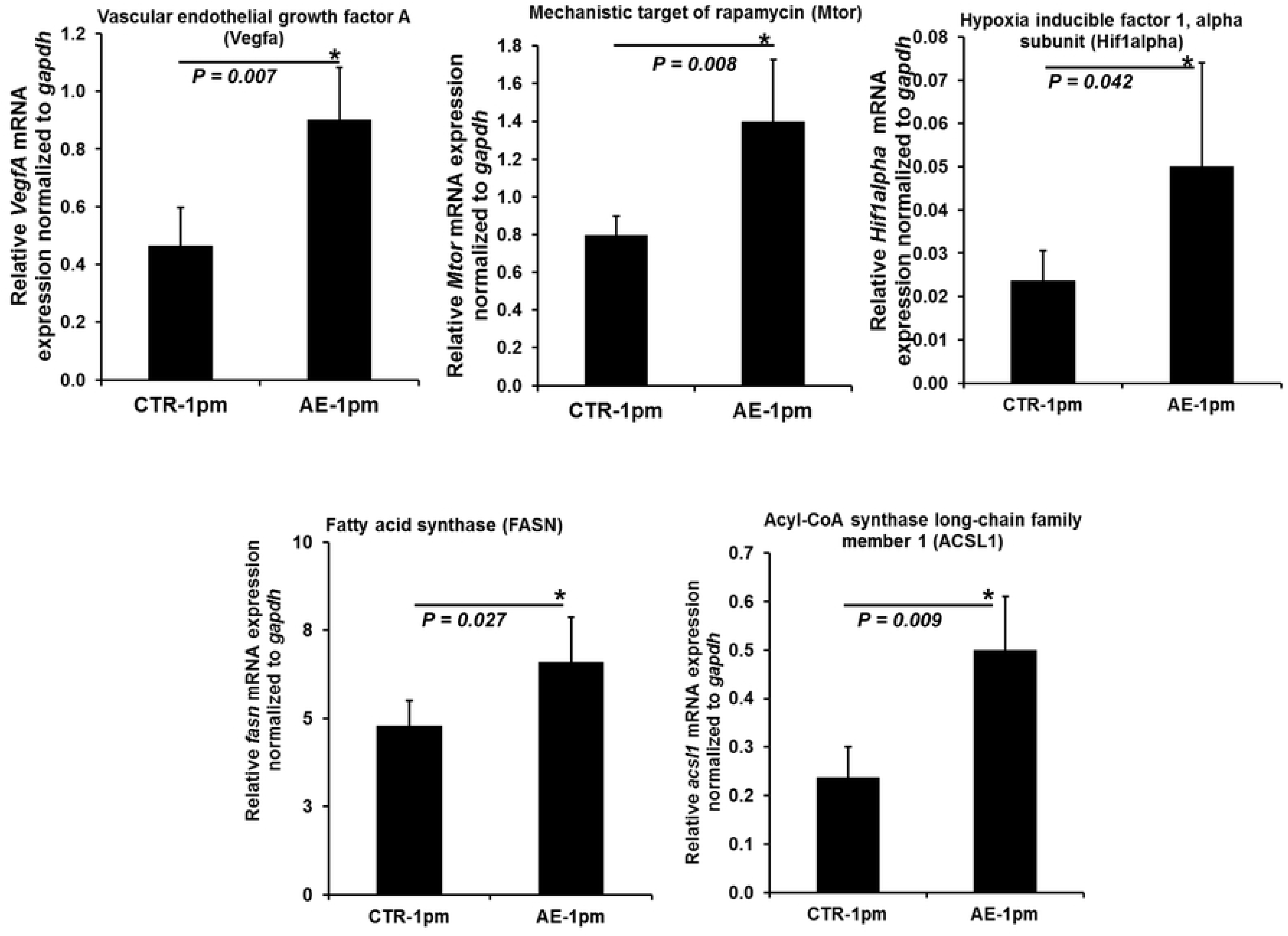
Relative mRNA level of angiogenesis- and lipid metabolism-related genes. Normalization was done using *gapdh* as endogenous reference. Bars represent median ± standard deviation. All qPCRs were carried-out using biological replicates (n = 5 mice from the infected group and n = 4 mice from uninfected group), with three technical replicates per sample.

## Discussion

In the present study we undertook NGS-based miRNA profiling in livers of mice at an early stage of primary AE. 28 miRNAs were identified that exhibited statistically significant dysregulated expression levels in the *E. multilocularis* infected liver.

Many reports have demonstrated that alterations in endogenous miRNA expression patterns correlated with various liver disorders including infectious- and non-infectious liver diseases [66]. Changes in miRNA profiles are also distincted between different liver injuries and were etiology-dependent [66]. This observation was consistent with our results since human AE affects the liver in 98% of cases and is considered as a chronic and progressive parasitic liver disease.

Previous studies have demonstrated changes in the liver transcriptome at early stages of *E. multilocularis* infection [49, 50], which could be very likely caused by the modified miRNAs expression found here, since miRNAs are key post-transcriptional regulators of gene expression [67]. For example, the expression level of the *Fasn* gene coding for fatty acid synthase was the most up-regulated at two month post-infection [50], and *Fasn* is a primary target of miR-15a-5p [68, 69] (down-regulated here), thus this confirmed the negative correlation between expression of a given miRNA and its target gene. However, studies on transcriptome characterization of both mRNA and miRNA during AE should be done concurrently, because comparisons between studies are not always feasible/reasonable due to experimental variables.

Many reports have shown that alterations of miRNAs expression patterns can be disease stage–dependent, with presence of miRNA signatures common to all clinical phases [70, 71]. In our study, we only examined miRNA expression profiles at the early stage of AE (one month post-infection), thus future investigations will be designed to measure the hepatic miRNA responses at later stages of the disease.

Human AE is clinically/symptomatically often compared to liver cancer, since metacestodes exhibit progressive growth of lesions and a capacity to spread to other organs by metastatis formation [5, 72]. Nineteen out of the 28 dysregulated miRNAs identified in our study have been reported to be also affected in hepatocellular carcinoma (HCC) which is the most common type of primary liver cancer in adults [73–81]. Moreover, miR-148a is a tumor suppressor and is down-regulated in HCC and in various cancers, and miR-148a plays crucial roles in migration, invasion and apoptosis of tumor cells [73].

In terms of immunity, both miR-148a-3p and miR-30a-5p modulate inflammation by repressing NF-κB signaling in macrophages and respective pro-inflammatory consequences [82, 83]. It was shown that the use of miRNAs mimicking miR-30a significantly down-regulates the expression levels of IL-1α, IL-6 and TNF-α mRNAs in LPS-stimulated macrophages [82]. In our study, miR-148a-3p and miR-30a-5p were dramatically down-regulated, which is consistent with previous reports showing that in mice, primary AE is characterized by an initial acute inflammatory Th1 immune response which then gradually shifts to a Th2 response due to an immunomodulatory down-regulation effect presumably mediated by the *E. multilocularis* larvae [8, 84].

In terms of lipid metabolism, several studies have shown that miR-148a (5p and 3p) is directly involved in controlling cholesterol and triglyceride homeostasis and circulating lipoprotein levels [85–87]. Deficiency or low levels of miR-148a (5p and 3p) promotes hepatic lipid metabolism. Furthermore, miR-148a knockout mice suffer from fatty liver disease (hepatic steatosis) that is associated with accumulation of triglyceride fat in large vacuoles within liver cells and with significantly increased expression of genes involved in lipogenesis and fatty acid uptake [88]. In our study, both miR-148a-5p and miR-148a-3p were down-regulated, suggesting that infection with *E. multilocularis* may lead to increased lipid synthesis in the liver of the host. This hypothesis is strongly supported by the significantly high mRNA levels of the two lipogenic enzymes FASN and ACSL1 in livers of *E. multilocularis*-infected mice when compared to controls. Fatty acid synthase (FASN) catalyzes the last step in fatty acid biosynthesis, and thus, its expression level is a major factor in determining the maximal hepatic capacity to generate fatty acids by *de novo* lipogenesis [89]. In the liver, ACSL1 is the predominant isoform of the long-chain acyl-CoA synthetase family and it accounts for 50% of total hepatic ACSL activity [90]. Over expression of ACSL1 results in increased proportion of oleic acid in diacylglycerol (DAG) and phospholipids (PLs), and promotes synthesis of triglyceride (TG) from free fatty acids and its accumulation in hepatoma cells [91].

The liver-specific miRNA-122 may account for up to 70% of the total miRNA pool and its endogenous and circulating levels are consistently affected in close association with liver diseases [92]. In hepatocellular carcinoma, miR-122 was shown to be down-regulated in nearly 70% of cases [93], whereas in our study, primary AE led to increased miR-122-5p expression. The microRNA miR-122 plays an essential role in the regulation of cholesterol and fatty acid (FAs) metabolism [92]; in mice, inhibition of miR-122 by *in vivo* antisense targeting resulted in a decrease in hepatic FAs and cholesterol synthesis rates [94]. Thus, despite the great resemblance we found here between HCC and murine primary AE in term of miRNA dysregulation pattern, there is an important difference with respect to miR-122. This difference can be explained by the fact that FAs have different implications in various infectious and non-infectious liver diseases. In fact, in contrast to tumor cells, *E. multilocularis* larvae are unable to synthesize FAs, cholesterol and other sterols *de novo*, thus they must scavenge these components from their host [95], and mobilization of host lipid resources to the parasite may imply stimulation of fatty acid biosynthesis [96].

Interestingly, our pathway enrichment analysis of down-regulated miRNAs revealed a clear enrichment for genes positively regulating angiogenesis (VEGFA/VEGFR2 signaling pathway) and its related functions (cell proliferation, cell migration, cell survival, as well as VEGFR2 endocytosis and recycling) [97]. Angiogenesis is a complex multistep process that leads to formation of new blood vessels from existing vessels. This process allows tumor cells to be supplied by oxygen and nutrients and dispose of waste products. Among the pro-angiogenic endogenous molecules, vascular endothelial growth factor A (VEGFA) is a major regulator of blood vessel formation and it binds to VEGF receptor 2 (VEGFR2) expressed by endothelial cells [97, 98]. In HCC and in response to hypoxic conditions, transcription of *vegfa* is promoted by hypoxia-inducible factor 1-alpha (HIF1-α) [99]. Moreover, HIF-1α has been shown to be expressed significantly higher in the actively multiplying infiltrative region of the AE liver lesions in comparison to the hepatic parenchyma of Wistar rats [100]. Herein, relative mRNA levels of *vegfa* and *hif1-α* were significantly higher in *E. multilocularis* infected liver tissues, which still emphasized the similarity between HCC and hepatic AE. Both *vegfa* and *hif1-α* are experimentally validated targets of mmu-miR-15a-5p, mmu-miR-126a-5p and mmu-miR-101a/b-3p [101–103] (down-regulated miRNAs in this study).

In pathological situations, angiogenesis is driven by an immune reaction that includes the secretion of proinflammatory cytokines particularly IL-1β, TNF-α [104] and IL-6 [105]. Secretion of IL-1β, TNF-α and IL-6 is hallmark of the early inflammatory phase of AE observed in mice [84]. The mechanistic target of rapamycin, mTOR, which is targeted by the down-regulated mmu-miR-101a/b-3p mmu-miR-15a-5p, has recently emerged as a regulator linking inflammation to angiogenesis trough TNFα/IKKβ signaling pathway [106]. Activation of mTOR leads to the production and release of the extracellular matrix-degrading and remodeling enzymes such as matrix metalloproteinase 2 and 9 (MMP2, MMP9) and urokinase plasminogen activator (uPA) facilitating thus the migration of proliferating endothelial cells to form new vessels [107]. In this study, the relative expression of *mtor* was also significantly higher in *E. multilocularis* infected liver tissues. Overall, the findings reported herein provided a strong molecular evidence for an *E. multilocularis* induced angiogenesis at early stage of the infection. This phenomenon is at least partially mediated via downregulation of specific miRNAs such as mmu-miR-15a-5p, mmu-miR-126a-5p and mmu-miR-101a/b-3p. From a mechanistic point of view and inferring to gene expression and immunological studies on AE, the *E. multilocularis* induced angiogenesis might be a consequence of (i) the early immune host reaction with over-expression of inflammation-related cytokines [84], chemokines [50] and metalloproteinases (MMPs) [108] and (ii) the progressive increase of tissue hypoxia caused by anatomical modifications in the compressed hepatic areas surrounding the continuously growing metacestodes. Overall, the role of angiogenesis during AE remains largely unknown, despite the fact that this phenomenon occurs during infection with *E. multilocularis* [8, 109]. Although the metacestode tissue itself is not vascularized, issues such as the role of periparasitic angiogenesis favouring periparasitic nutrition, a provision and putatively metastasis formation at late AE stage need to be addressed. It was recently shown that in human AE, the angiographic vascularity of *E. multilocularis* lesions is detectable and measurable using dual-energy computer tomography [109]. In view of the resemblance between liver cancer and AE on one hand and the important role of angiogenesis in the development and growth of metastasis on the other hand, future studies on understanding the impact of the VEGF/VEGFR pathway during AE are needed; such investigations may open a new window for the assessment of anti-angiogenic therapeutics for human AE [110].

In this study, we also examined arm selection preferences (5p or 3p) in miRNA pairs of normal and *E. multilocularis* infected liver tissues. In fact, it was widely reported that the double stranded pre-miRNAs produced only one mature functional miRNA at one arm, either 5p or 3p. This selection relied largely on the thermodynamic stability of the strands as a chief factor in determining which arm of the duplex will be incorporated in the RISC complex as functional miRNA. Recent evidence revealed that many pre-miRNAs may yield two mature products deriving from both arms (5p and 3p) with different selection rates [111–114].

Furthermore, it was shown that in response to a pathological condition e.g. cancer [115, 116] and infection [117], the 5p- and 3p selection preference of some pre-miRNAs was altered. So this so-called “arm switching” phenomenon is now recognized as a miRNA post-transcriptional regulatory mechanism [118]. In our study, we found nine miRNAs showing significant differences in 5p- and 3p-arm selection preference between normal and infected liver tissue. Arm switching may be explained by the fact that the miR-5p and miR-3p resulting from the same pre-miRNA may act on different mRNA targets, thus they might be subjected to inverse regulation [119].

In conclusion, our data highlighted the role of hepatic microRNAs as molecular regulators that control early host reaction to *E. multilocularis* infection. We showed that liver miRNA transcriptome became significantly altered during the early stage of primary AE. This change was characterized by a high selective regulation of miRNA molecules which are involved in metabolic and cellular processes beneficial for *E. multilocularis* larval growth and proliferation, in particular angiogenesis and lipid synthesis.

Although the early phase of primary AE is crucial for understanding the first key molecular events that contribute to the establishment of *E. multilocularis* larvae, this stage is not the ideal target for treatment since the vast majority of AE-patients are diagnosed at advanced stages of the disease. Consequently, to that, future work on the expression changes of miRNAs during AE progression is urgently required with the prospective goal to first identify miRNAs signatures associated with the disease progression, and second evaluate microRNA-based therapeutics.

For the most significantly dysregulated hepatic miRNAs, future studies will also have to determine whether those molecules could be detected in the serum/plasma and if yes whether they are subjected to the same type of regulation. Such investigations are necessary for assessing microRNAs as new class of biomarkers, which hold a great potential to improve time to diagnosis and predict response to therapy.

## Acknowledgements

The authors gratefully acknowledge Dr. Marcella Cucher for critical reading of the manuscript (Instituto de Investigaciones en Microbiología y Parasitología Médica, Buenos Aires, Argentina).

## Competing Interests

Sebastian Strempel is employee of Microsynth AG.

## Supporting information

**Supplementary Fig 1. Down-regulated microRNA-target network.** This network represents regulatory relationships between down-regulated miRNAs in alveolar echinococcois and their target genes. Blue square: miRNAs and red dots: target genes. The 25 genes involved in VEGFA-VEGFR2 pathway are highlighted in green. The Network can be reproduced by entering the set of down-regulated miRNAs online in http://www.mirnet.ca/.

**Supplementary Fig 2. Up-regulated microRNA-target network**. In liver of *E. multilocularis*-infected mice, 9 miRNAs were significantly overexpressed as compared to uninfected controls. Target genes of seven up-regulated miRNAs were predicted. This network represents regulatory relationships between up-regulated miRNAs and their target genes. Two miRNAs (mmu-mir-1839-5p and mmu-mir-28a-5p) were not connected neither to the main tree nor to each other, thus they are not presented here. Twenty-six genes are common between two microRNAs (dark yellow circles). The seven genes involved in heme biosynthesis pathway are highlighted in green.

## References

1. Eckert J, Deplazes P. Biological, Epidemiological, and Clinical Aspects of Echinococcosis, a Zoonosis of Increasing Concern. Clin Microbiol Rev. 2004;17: 107–135. doi:10.1128/CMR.17.1.107-135.2004

2. Otero-Abad B, Rüegg SR, Hegglin D, Deplazes P, Torgerson PR. Mathematical modelling of *Echinococcus multilocularis* abundance in foxes in Zurich, Switzerland. Parasit Vectors. 2017;10: 21. doi:10.1186/s13071-016-1951-1

3. Bardonnet K, Vuitton DA, Grenouillet F, Mantion GA, Delabrousse E, Blagosklonov O, et al. 30-yr course and favorable outcome of alveolar echinococcosis despite multiple metastatic organ involvement in a non-immune suppressed patient. Ann Clin Microbiol Antimicrob. 2013;12: 1. doi:10.1186/1476-0711-12-1

4. Brunetti E, Kern P, Vuitton DA, Writing Panel for the WHO-IWGE. Expert consensus for the diagnosis and treatment of cystic and alveolar echinococcosis in humans. Acta Trop. 2010;114: 1–16. doi:10.1016/j.actatropica.2009.11.001

5. Caire Nail L, Rodríguez Reimundes E, Weibel Galluzzo C, Lebowitz D, Ibrahim YL, Lobrinus JA, et al. Disseminated alveolar echinococcosis resembling metastatic malignancy: a case report. J Med Case Rep. 2017;11: 113. doi:10.1186/s13256-017-1279-2

6. Tappe D, Weise D, Ziegler U, Müller A, Müllges W, Stich A. Brain and lung metastasis of alveolar echinococcosis in a refugee from a hyperendemic area. J Med Microbiol. 2008;57: 1420–1423. doi:10.1099/jmm.0.2008/002816-0

7. Mejri N, Hemphill A, Gottstein B. Triggering and modulation of the host-parasite interplay by *Echinococcus multilocularis*: a review. Parasitology. 2010;137: 557–568. doi:10.1017/S0031182009991533

8. Vuitton DA, Gottstein B. *Echinococcus multilocularis* and Its Intermediate Host: A Model of Parasite-Host Interplay. In: BioMed Research International [Internet]. 2010 [cited 20 Dec 2017]. doi:10.1155/2010/923193

9. Torgerson PR, Schweiger A, Deplazes P, Pohar M, Reichen J, Ammann RW, et al. Alveolar echinococcosis: from a deadly disease to a well-controlled infection. Relative survival and economic analysis in Switzerland over the last 35 years. J Hepatol. 2008;49: 72–77. doi:10.1016/j.jhep.2008.03.023

10. Guidelines for treatment of cystic and alveolar echinococcosis in humans. WHO Informal Working Group on Echinococcosis. Bull World Health Organ. 1996;74: 231–242.

11. Hemphill A, Stadelmann B, Rufener R, Spiliotis M, Boubaker G, Müller J, et al. Treatment of echinococcosis: albendazole and mebendazole--what else? Parasite. 2014;21: 70. doi:10.1051/parasite/2014073

12. Lee RC, Feinbaum RL, Ambros V. The C. elegans heterochronic gene lin-4 encodes small RNAs with antisense complementarity to lin-14. Cell. 1993;75: 843–854.

13. Ha M, Kim VN. Regulation of microRNA biogenesis. Nat Rev Mol Cell Biol. 2014;15: 509–524. doi:10.1038/nrm3838

14. Kim VN. Small RNAs: classification, biogenesis, and function. Mol Cells. 2005;19: 1–15.

15. Hafner M, Landthaler M, Burger L, Khorshid M, Hausser J, Berninger P, et al. Transcriptome-wide identification of RNA-binding protein and microRNA target sites by PAR-CLIP. Cell. 2010;141: 129–141. doi:10.1016/j.cell.2010.03.009

16. Lytle JR, Yario TA, Steitz JA. Target mRNAs are repressed as efficiently by microRNA-binding sites in the 5’ UTR as in the 3’ UTR. Proc Natl Acad Sci USA. 2007;104: 9667–9672. doi:10.1073/pnas.0703820104

17. Huntzinger E, Izaurralde E. Gene silencing by microRNAs: contributions of translational repression and mRNA decay. Nat Rev Genet. 2011;12: 99–110. doi:10.1038/nrg2936

18. Bentwich I, Avniel A, Karov Y, Aharonov R, Gilad S, Barad O, et al. Identification of hundreds of conserved and nonconserved human microRNAs. Nat Genet. 2005;37: 766–770. doi:10.1038/ng1590

19. Flicek P, Amode MR, Barrell D, Beal K, Billis K, Brent S, et al. Ensembl 2014. Nucleic Acids Res. 2014;42: D749–755. doi:10.1093/nar/gkt1196

20. Londin E, Loher P, Telonis AG, Quann K, Clark P, Jing Y, et al. Analysis of 13 cell types reveals evidence for the expression of numerous novel primate- and tissue-specific microRNAs. Proc Natl Acad Sci USA. 2015;112: E1106–1115. doi:10.1073/pnas.1420955112

21. Friedman RC, Farh KK-H, Burge CB, Bartel DP. Most mammalian mRNAs are conserved targets of microRNAs. Genome Res. 2009;19: 92–105. doi:10.1101/gr.082701.108

22. Lewis BP, Burge CB, Bartel DP. Conserved seed pairing, often flanked by adenosines, indicates that thousands of human genes are microRNA targets. Cell. 2005;120: 15–20. doi:10.1016/j.cell.2004.12.035

23. Dumortier O, Hinault C, Van Obberghen E. MicroRNAs and metabolism crosstalk in energy homeostasis. Cell Metab. 2013;18: 312–324. doi:10.1016/j.cmet.2013.06.004

24. Mehta A, Baltimore D. MicroRNAs as regulatory elements in immune system logic. Nat Rev Immunol. 2016;16: 279–294. doi:10.1038/nri.2016.40

25. Song G, Sharma AD, Roll GR, Ng R, Lee AY, Blelloch RH, et al. MicroRNAs control hepatocyte proliferation during liver regeneration. Hepatology. 2010;51: 1735–1743. doi:10.1002/hep.23547

26. Das J, Podder S, Ghosh TC. Insights into the miRNA regulations in human disease genes. BMC Genomics. 2014;15: 1010. doi:10.1186/1471-2164-15-1010

27. Lin S, Gregory RI. MicroRNA biogenesis pathways in cancer. Nat Rev Cancer. 2015;15: 321–333. doi:10.1038/nrc3932

28. Kota J, Chivukula RR, O’Donnell KA, Wentzel EA, Montgomery CL, Hwang H-W, et al. Therapeutic microRNA delivery suppresses tumorigenesis in a murine liver cancer model. Cell. 2009;137: 1005–1017. doi:10.1016/j.cell.2009.04.021

29. Ling H, Fabbri M, Calin GA. MicroRNAs and other non-coding RNAs as targets for anticancer drug development. Nat Rev Drug Discov. 2013;12: 847–865. doi:10.1038/nrd4140

30. Zhang J, Chong CCN, Chen GG, Lai PBS. A Seven-microRNA Expression Signature Predicts Survival in Hepatocellular Carcinoma. PLoS ONE. 2015;10: e0128628. doi:10.1371/journal.pone.0128628

31. Ji J, Shi J, Budhu A, Yu Z, Forgues M, Roessler S, et al. MicroRNA expression, survival, and response to interferon in liver cancer. N Engl J Med. 2009;361: 1437–1447. doi:10.1056/NEJMoa0901282

32. Jopling CL, Yi M, Lancaster AM, Lemon SM, Sarnow P. Modulation of hepatitis C virus RNA abundance by a liver-specific MicroRNA. Science. 2005;309: 1577–1581. doi:10.1126/science.1113329

33. Britton C, Winter AD, Marks ND, Gu H, McNeilly TN, Gillan V, et al. Application of small RNA technology for improved control of parasitic helminths. Vet Parasitol. 2015;212: 47–53. doi:10.1016/j.vetpar.2015.06.003

34. Cabantous S, Hou X, Louis L, He H, Mariani O, Sastre X, et al. Evidence for an important role of host microRNAs in regulating hepatic fibrosis in humans infected with Schistosoma japonicum. Int J Parasitol. 2017;47: 823–830. doi:10.1016/j.ijpara.2017.05.007

35. Cai P, Gobert GN, McManus DP. MicroRNAs in Parasitic Helminthiases: Current Status and Future Perspectives. Trends Parasitol. 2016;32: 71–86. doi:10.1016/j.pt.2015.09.003

36. Han S, Tang Q, Lu X, Chen R, Li Y, Shu J, et al. Dysregulation of hepatic microRNA expression profiles with Clonorchis sinensis infection. BMC Infect Dis. 2016;16: 724. doi:10.1186/s12879-016-2058-1

37. Arora N, Tripathi S, Singh AK, Mondal P, Mishra A, Prasad A. Micromanagement of Immune System: Role of miRNAs in Helminthic Infections. Front Microbiol. 2017;8: 586. doi:10.3389/fmicb.2017.00586

38. He X, Sai X, Chen C, Zhang Y, Xu X, Zhang D, et al. Host serum miR-223 is a potential new biomarker for Schistosoma japonicum infection and the response to chemotherapy. Parasit Vectors. 2013;6: 272. doi:10.1186/1756-3305-6-272

39. Guo X, Zheng Y. Expression profiling of circulating miRNAs in mouse serum in response to *Echinococcus multilocularis* infection. Parasitology. 2017;144: 1079–1087. doi:10.1017/S0031182017000300

40. Jin X, Guo X, Zhu D, Ayaz M, Zheng Y. miRNA profiling in the mice in response to *Echinococcus multilocularis* infection. Acta Trop. 2017;166: 39–44. doi:10.1016/j.actatropica.2016.10.024

41. Jiang S, Li X, Wang X, Ban Q, Hui W, Jia B. MicroRNA profiling of the intestinal tissue of Kazakh sheep after experimental *Echinococcus granulosus* infection, using a high-throughput approach. Parasite. 2016;23: 23. doi:10.1051/parasite/2016023

42. Christopher AF, Kaur RP, Kaur G, Kaur A, Gupta V, Bansal P. MicroRNA therapeutics: Discovering novel targets and developing specific therapy. Perspect Clin Res. 2016;7: 68–74. doi:10.4103/2229-3485.179431

43. Ancarola ME, Marcilla A, Herz M, Macchiaroli N, Pérez M, Asurmendi S, et al. Cestode parasites release extracellular vesicles with microRNAs and immunodiagnostic protein cargo. Int J Parasitol. 2017;47: 675–686. doi:10.1016/j.ijpara.2017.05.003

44. Macchiaroli N, Maldonado LL, Zarowiecki M, Cucher M, Gismondi MI, Kamenetzky L, et al. Genome-wide identification of microRNA targets in the neglected disease pathogens of the genus *Echinococcus*. Mol Biochem Parasitol. 2017;214: 91–100. doi:10.1016/j.molbiopara.2017.04.001

45. Cucher M, Macchiaroli N, Kamenetzky L, Maldonado L, Brehm K, Rosenzvit MC. High-throughput characterization of *Echinococcus* spp. metacestode miRNomes. Int J Parasitol. 2015;45: 253–267. doi:10.1016/j.ijpara.2014.12.003

46. Britton C, Winter AD, Gillan V, Devaney E. microRNAs of parasitic helminths - Identification, characterization and potential as drug targets. Int J Parasitol Drugs Drug Resist. 2014;4: 85–94. doi:10.1016/j.ijpddr.2014.03.001

47. Kepron C, Novak M, Blackburn BJ. Effect of *Echinococcus multilocularis* on the origin of acetyl-coA entering the tricarboxylic acid cycle in host liver. J Helminthol. 2002;76: 31–36.

48. Novak M, Modha A, Blackburn BJ. Metabolic alterations in organs of Meriones unguiculatus infected with *Echinococcus multilocularis*. Comp Biochem Physiol, B. 1993;105: 517–521.

49. Gottstein B, Wittwer M, Schild M, Merli M, Leib SL, Müller N, et al. Hepatic Gene Expression Profile in Mice Perorally Infected with *Echinococcus multilocularis* Eggs. PLoS ONE. 2010;5: e9779. doi:10.1371/journal.pone.0009779

50. Lin R, Lü G, Wang J, Zhang C, Xie W, Lu X, et al. Time course of gene expression profiling in the liver of experimental mice infected with *Echinococcus multilocularis*. PLoS ONE. 2011;6: e14557. doi:10.1371/journal.pone.0014557

51. Deplazes P, Grimm F, Sydler T, Tanner I, Kapel CMO. Experimental alveolar echinococcosis in pigs, lesion development and serological follow up. Vet Parasitol. 2005;130: 213–222. doi:10.1016/j.vetpar.2005.03.034

52. Edgar RC. Search and clustering orders of magnitude faster than BLAST. Bioinformatics. 2010;26: 2460–2461. doi:10.1093/bioinformatics/btq461

53. Kozomara A, Griffiths-Jones S. miRBase: annotating high confidence microRNAs using deep sequencing data. Nucleic Acids Res. 2014;42: D68–D73. doi:10.1093/nar/gkt1181

54. Griffiths-Jones S, Saini HK, van Dongen S, Enright AJ. miRBase: tools for microRNA genomics. Nucleic Acids Res. 2008;36: D154–D158. doi:10.1093/nar/gkm952

55. Nawrocki EP, Burge SW, Bateman A, Daub J, Eberhardt RY, Eddy SR, et al. Rfam 12.0: updates to the RNA families database. Nucleic Acids Res. 2015;43: D130–137. doi:10.1093/nar/gku1063

56. Dobin A, Davis CA, Schlesinger F, Drenkow J, Zaleski C, Jha S, et al. STAR: ultrafast universal RNA-seq aligner. Bioinformatics. 2013;29: 15–21. doi:10.1093/bioinformatics/bts635

57. Anders S, Pyl PT, Huber W. HTSeq—a Python framework to work with high-throughput sequencing data. Bioinformatics. 2015;31: 166–169. doi:10.1093/bioinformatics/btu638

58. Love MI, Huber W, Anders S. Moderated estimation of fold change and dispersion for RNA-seq data with DESeq2. Genome Biol. 2014;15: 550. doi:10.1186/s13059-014-0550-8

59. Robinson MD, McCarthy DJ, Smyth GK. edgeR: a Bioconductor package for differential expression analysis of digital gene expression data. Bioinformatics. 2010;26: 139–140. doi:10.1093/bioinformatics/btp616

60. Ringnér M. What is principal component analysis? Nat Biotechnol. 2008;26: 303–304. doi:10.1038/nbt0308-303

61. Chen C, Ridzon DA, Broomer AJ, Zhou Z, Lee DH, Nguyen JT, et al. Real-time quantification of microRNAs by stem-loop RT-PCR. Nucleic Acids Res. 2005;33: e179. doi:10.1093/nar/gni178

62. Matoušková P, Bártíková H, Boušová I, Hanušová V, Szotáková B, Skálová L. Reference genes for real-time PCR quantification of messenger RNAs and microRNAs in mouse model of obesity. PLoS ONE. 2014;9: e86033. doi:10.1371/journal.pone.0086033

63. Fan Y, Siklenka K, Arora SK, Ribeiro P, Kimmins S, Xia J. miRNet - dissecting miRNA-target interactions and functional associations through network-based visual analysis. Nucleic Acids Res. 2016;44: W135–141. doi:10.1093/nar/gkw288

64. D’Eustachio P. Reactome knowledgebase of human biological pathways and processes. Methods Mol Biol. 2011;694: 49–61. doi:10.1007/978-1-60761-977-2_4

65. Kanehisa M, Araki M, Goto S, Hattori M, Hirakawa M, Itoh M, et al. KEGG for linking genomes to life and the environment. Nucleic Acids Res. 2008;36: D480–484. doi:10.1093/nar/gkm882

66. Szabo G, Bala S. MicroRNAs in liver disease. Nat Rev Gastroenterol Hepatol. 2013;10: 542–552. doi:10.1038/nrgastro.2013.87

67. Bentwich I, Avniel A, Karov Y, Aharonov R, Gilad S, Barad O, et al. Identification of hundreds of conserved and nonconserved human microRNAs. Nat Genet. 2005;37: 766–770. doi:10.1038/ng1590

68. Chen Z, Qiu H, Ma L, Luo J, Sun S, Kang K, et al. miR-30e-5p and miR-15a Synergistically Regulate Fatty Acid Metabolism in Goat Mammary Epithelial Cells via LRP6 and YAP1. Int J Mol Sci. 2016;17. doi:10.3390/ijms17111909

69. Wang J, Zhang X, Shi J, Cao P, Wan M, Zhang Q, et al. Fatty acid synthase is a primary target of MiR-15a and MiR-16-1 in breast cancer. Oncotarget. 2016;7: 78566–78576. doi:10.18632/oncotarget.12479

70. Xing T, Xu H, Yu W, Wang B, Zhang J. Expression profile and clinical significance of miRNAs at different stages of chronic hepatitis B virus infection. Int J Clin Exp Med. 2015;8: 5611–5620.

71. Bandyopadhyay S, Long ME, Allen L-AH. Differential expression of microRNAs in Francisella tularensis-infected human macrophages: miR-155-dependent downregulation of MyD88 inhibits the inflammatory response. PLoS ONE. 2014;9: e109525. doi:10.1371/journal.pone.0109525

72. Bresson-Hadni S, Miguet J-P, Mantion G, Giraudoux P, Vuitton D-A. [Alveolar echinococcosis: a disease comparable to a slow growing cancer]. Bull Acad Natl Med. 2008;192: 1131–1138; discussion 1139.

73. Li Y, Deng X, Zeng X, Peng X. The Role of Mir-148a in Cancer. Journal of Cancer. 2016;7: 1233–1241. doi:10.7150/jca.14616

74. Liu B, Sun T, Wu G, Shang-Guan H, Jiang Z-J, Zhang J-R, et al. MiR-15a suppresses hepatocarcinoma cell migration and invasion by directly targeting cMyb. Am J Transl Res. 2017;9: 520–532.

75. Raitoharju E, Seppälä I, Lyytikäinen L-P, Viikari J, Ala-Korpela M, Soininen P, et al. Blood hsa-miR-122-5p and hsa-miR-885-5p levels associate with fatty liver and related lipoprotein metabolism-The Young Finns Study. Sci Rep. 2016;6: 38262. doi:10.1038/srep38262

76. Su X, Wang H, Ge W, Yang M, Hou J, Chen T, et al. An In Vivo Method to Identify microRNA Targets Not Predicted by Computation Algorithms: p21 Targeting by miR-92a in Cancer. Cancer Res. 2015;75: 2875–2885. doi:10.1158/0008-5472.CAN-14-2218

77. Sun J, Lu H, Wang X, Jin H. MicroRNAs in hepatocellular carcinoma: regulation, function, and clinical implications. ScientificWorldJournal. 2013;2013: 924206. doi:10.1155/2013/924206

78. Thurnherr T, Mah W-C, Lei Z, Jin Y, Rozen SG, Lee CG. Differentially Expressed miRNAs in Hepatocellular Carcinoma Target Genes in the Genetic Information Processing and Metabolism Pathways. Sci Rep. 2016;6: 20065. doi:10.1038/srep20065

79. Xu L, Beckebaum S, Iacob S, Wu G, Kaiser GM, Radtke A, et al. MicroRNA-101 inhibits human hepatocellular carcinoma progression through EZH2 downregulation and increased cytostatic drug sensitivity. J Hepatol. 2014;60: 590–598. doi:10.1016/j.jhep.2013.10.028

80. Zhang J, Yang Y, Yang T, Liu Y, Li A, Fu S, et al. microRNA-22, downregulated in hepatocellular carcinoma and correlated with prognosis, suppresses cell proliferation and tumourigenicity. Br J Cancer. 2010;103: 1215–1220. doi:10.1038/sj.bjc.6605895

81. Zhuang L, Wang X, Wang Z, Ma X, Han B, Zou H, et al. MicroRNA-23b functions as an oncogene and activates AKT/GSK3β/β-catenin signaling by targeting ST7L in hepatocellular carcinoma. Cell Death Dis. 2017;8: e2804. doi:10.1038/cddis.2017.216

82. Jiang X, Xu C, Lei F, Liao M, Wang W, Xu N, et al. MiR-30a targets IL-1α and regulates islet functions as an inflammation buffer and response factor. Sci Rep. 2017;7: 5270. doi:10.1038/s41598-017-05560-1

83. Patel V, Carrion K, Hollands A, Hinton A, Gallegos T, Dyo J, et al. The stretch responsive microRNA miR-148a-3p is a novel repressor of IKBKB, NF-κB signaling, and inflammatory gene expression in human aortic valve cells. FASEB J. 2015;29: 1859–1868. doi:10.1096/fj.14-257808

84. Wang J, Lin R, Zhang W, Li L, Gottstein B, Blagosklonov O, et al. Transcriptional profiles of cytokine/chemokine factors of immune cell-homing to the parasitic lesions: a comprehensive one-year course study in the liver of E. multilocularis-infected mice. PLoS ONE. 2014;9: e91638. doi:10.1371/journal.pone.0091638

85. Wagschal A, Najafi-Shoushtari SH, Wang L, Goedeke L, Sinha S, deLemos AS, et al. Genome-wide identification of microRNAs regulating cholesterol and triglyceride homeostasis. Nat Med. 2015;21: 1290–1297. doi:10.1038/nm.3980

86. Goedeke L, Rotllan N, Canfrán-Duque A, Aranda JF, Ramírez CM, Araldi E, et al. MicroRNA-148a regulates LDL receptor and ABCA1 expression to control circulating lipoprotein levels. Nat Med. 2015;21: 1280–1289. doi:10.1038/nm.3949

87. Goedeke L, Wagschal A, Fernández-Hernando C, Näär AM. miRNA regulation of LDL-cholesterol metabolism. Biochim Biophys Acta. 2016;1861: 2047–2052. doi:10.1016/j.bbalip.2016.03.007

88. Cheng L, Zhu Y, Han H, Zhang Q, Cui K, Shen H, et al. MicroRNA-148a deficiency promotes hepatic lipid metabolism and hepatocarcinogenesis in mice. Cell Death Dis. 2017;8: e2916. doi:10.1038/cddis.2017.309

89. Dorn C, Riener M-O, Kirovski G, Saugspier M, Steib K, Weiss TS, et al. Expression of fatty acid synthase in nonalcoholic fatty liver disease. Int J Clin Exp Pathol. 2010;3: 505–514.

90. Li LO, Ellis JM, Paich HA, Wang S, Gong N, Altshuller G, et al. Liver-specific loss of long chain acyl-CoA synthetase-1 decreases triacylglycerol synthesis and beta-oxidation and alters phospholipid fatty acid composition. J Biol Chem. 2009;284: 27816–27826. doi:10.1074/jbc.M109.022467

91. Yan S, Yang X-F, Liu H-L, Fu N, Ouyang Y, Qing K. Long-chain acyl-CoA synthetase in fatty acid metabolism involved in liver and other diseases: An update. World J Gastroenterol. 2015;21: 3492–3498. doi:10.3748/wjg.v21.i12.3492

92. Jopling C. Liver-specific microRNA-122. RNA Biol. 2012;9: 137–142. doi:10.4161/rna.18827

93. Gupta P, Cairns MJ, Saksena NK. Regulation of gene expression by microRNA in HCV infection and HCV–mediated hepatocellular carcinoma. Virol J. 2014;11: 64. doi:10.1186/1743-422X-11-64

94. Esau C, Davis S, Murray SF, Yu XX, Pandey SK, Pear M, et al. miR-122 regulation of lipid metabolism revealed by in vivo antisense targeting. Cell Metab. 2006;3: 87–98. doi:10.1016/j.cmet.2006.01.005

95. Alvite G, Esteves A. Lipid binding proteins from parasitic platyhelminthes. Front Physiol. 2012;3. doi:10.3389/fphys.2012.00363

96. Kepron C, Schoen J, Novak M, J Blackburn B. NMR study of lipid changes in organs of jirds infected with *Echinococcus multilocularis*. Comparative Biochemistry and Physiology Part B: Biochemistry and Molecular Biology. 1999;124: 347–353. doi:10.1016/S0305-0491(99)00126-1

97. Abhinand CS, Raju R, Soumya SJ, Arya PS, Sudhakaran PR. VEGF-A/VEGFR2 signaling network in endothelial cells relevant to angiogenesis. J Cell Commun Signal. 2016;10: 347–354. doi:10.1007/s12079-016-0352-8

98. Bautch VL. VEGF-Directed Blood Vessel Patterning: From Cells to Organism. Cold Spring Harb Perspect Med. 2012;2. doi:10.1101/cshperspect.a006452

99. Chen C, Lou T. Hypoxia inducible factors in hepatocellular carcinoma. Oncotarget. 2017;8: 46691–46703. doi:10.18632/oncotarget.17358

100. Song T, Li H, Yang L, Lei Y, Yao L, Wen H. Expression of Hypoxia-Inducible Factor-1α in the Infiltrative Belt Surrounding Hepatic Alveolar Echinococcosis in Rats. J Parasitol. 2015;101: 369–373. doi:10.1645/14-685.1

101. Sun C-Y, She X-M, Qin Y, Chu Z-B, Chen L, Ai L-S, et al. miR-15a and miR-16 affect the angiogenesis of multiple myeloma by targeting VEGF. Carcinogenesis. 2013;34: 426–435. doi:10.1093/carcin/bgs333

102. Kong R, Ma Y, Feng J, Li S, Zhang W, Jiang J, et al. The crucial role of miR-126 on suppressing progression of esophageal cancer by targeting VEGF-A. Cell Mol Biol Lett. 2016;21. doi:10.1186/s11658-016-0004-2

103. Kim J-H, Lee K-S, Lee D-K, Kim J, Kwak S-N, Ha K-S, et al. Hypoxia-Responsive MicroRNA-101 Promotes Angiogenesis via Heme Oxygenase-1/Vascular Endothelial Growth Factor Axis by Targeting Cullin 3. Antioxid Redox Signal. 2014;21: 2469–2482. doi:10.1089/ars.2014.5856

104. TNF primes endothelial cells for angiogenic sprouting by inducing a tip cell phenotype. - PubMed - NCBI [Internet]. [cited 6 Jun 2019]. Available: https://www.ncbi.nlm.nih.gov/pubmed/18337563

105. Catar R, Witowski J, Zhu N, Lücht C, Derrac Soria A, Uceda Fernandez J, et al. IL-6 Trans-Signaling Links Inflammation with Angiogenesis in the Peritoneal Membrane. J Am Soc Nephrol. 2017;28: 1188–1199. doi:10.1681/ASN.2015101169

106. Lee D-F, Kuo H-P, Chen C-T, Hsu J-M, Chou C-K, Wei Y, et al. IKK beta suppression of TSC1 links inflammation and tumor angiogenesis via the mTOR pathway. Cell. 2007;130: 440–455. doi:10.1016/j.cell.2007.05.058

107. Conciatori F, Bazzichetto C, Falcone I, Pilotto S, Bria E, Cognetti F, et al. Role of mTOR Signaling in Tumor Microenvironment: An Overview. Int J Mol Sci. 2018;19. doi:10.3390/ijms19082453

108. Marco M, Baz A, Fernandez C, Gonzalez G, Hellman U, Salinas G, et al. A relevant enzyme in granulomatous reaction, active matrix metalloproteinase-9, found in bovine *Echinococcus granulosus* hydatid cyst wall and fluid. Parasitol Res. 2006;100: 131–139. doi:10.1007/s00436-006-0237-5

109. Jiang Y, Li J, Wang J, Xiao H, Li T, Liu H, et al. Assessment of Vascularity in Hepatic Alveolar Echinococcosis: Comparison of Quantified Dual-Energy CT with Histopathologic Parameters. PLoS ONE. 2016;11: e0149440. doi:10.1371/journal.pone.0149440

110. Dennis RD, Schubert U, Bauer C. Angiogenesis and parasitic helminth-associated neovascularization. Parasitology. 2011;138: 426–439. doi:10.1017/S0031182010001642

111. Kloosterman WP, Steiner FA, Berezikov E, de Bruijn E, van de Belt J, Verheul M, et al. Cloning and expression of new microRNAs from zebrafish. Nucleic Acids Res. 2006;34: 2558–2569. doi:10.1093/nar/gkl278

112. Ruby JG, Stark A, Johnston WK, Kellis M, Bartel DP, Lai EC. Evolution, biogenesis, expression, and target predictions of a substantially expanded set of Drosophila microRNAs. Genome Res. 2007;17: 1850–1864. doi:10.1101/gr.6597907

113. Okamura K, Phillips MD, Tyler DM, Duan H, Chou Y, Lai EC. The regulatory activity of microRNA* species has substantial influence on microRNA and 3’ UTR evolution. Nat Struct Mol Biol. 2008;15: 354–363. doi:10.1038/nsmb.1409

114. Tsai K-W, Leung C-M, Lo Y-H, Chen T-W, Chan W-C, Yu S-Y, et al. Arm Selection Preference of MicroRNA-193a Varies in Breast Cancer. Sci Rep. 2016;6: 28176. doi:10.1038/srep28176

115. Li S-C, Liao Y-L, Ho M-R, Tsai K-W, Lai C-H, Lin W. miRNA arm selection and isomiR distribution in gastric cancer. BMC Genomics. 2012;13 Suppl 1: S13. doi:10.1186/1471-2164-13-S1-S13

116. Li S-C, Tsai K-W, Pan H-W, Jeng Y-M, Ho M-R, Li W-H. MicroRNA 3’ end nucleotide modification patterns and arm selection preference in liver tissues. BMC Syst Biol. 2012;6 Suppl 2: S14. doi:10.1186/1752-0509-6-S2-S14

117. Siddle KJ, Tailleux L, Deschamps M, Loh Y-HE, Deluen C, Gicquel B, et al. bacterial infection drives the expression dynamics of microRNAs and their isomiRs. PLoS Genet. 2015;11: e1005064. doi:10.1371/journal.pgen.1005064

118. Guo L, Yu J, Yu H, Zhao Y, Chen S, Xu C, et al. Evolutionary and expression analysis of miR-#-5p and miR-#-3p at the miRNAs/isomiRs levels. Biomed Res Int. 2015;2015: 168358. doi:10.1155/2015/168358

119. Choo KB, Soon YL, Nguyen PNN, Hiew MSY, Huang C-J. MicroRNA-5p and -3p co-expression and cross-targeting in colon cancer cells. J Biomed Sci. 2014;21: 95. doi:10.1186/s12929-014-0095-x

120. Chen H, Tian Y. MiR-15a-5p regulates viability and matrix degradation of human osteoarthritis chondrocytes via targeting VEGFA. Biosci Trends. 2017;10: 482–488. doi:10.5582/bst.2016.01187

121. Liu B, Peng X-C, Zheng X-L, Wang J, Qin Y-W. MiR-126 restoration down-regulate VEGF and inhibit the growth of lung cancer cell lines in vitro and in vivo. Lung Cancer. 2009;66: 169–175. doi:10.1016/j.lungcan.2009.01.010

122. Wang C-Z, Deng F, Li H, Wang D-D, Zhang W, Ding L, et al. MiR-101: a potential therapeutic target of cancers. : 12.

123. Zhang Y, Bo H, Wang H-Y, Chang A, Zheng XFS. Emerging Role of MicroRNAs in mTOR Signaling. Cell Mol Life Sci. 2017;74: 2613–2625. doi:10.1007/s00018-017-2485-1

124. Yang J, Liu R, Deng Y, Qian J, Lu Z, Wang Y, et al. MiR-15a/16 deficiency enhances anti-tumor immunity of glioma-infiltrating CD8+ T cells through targeting mTOR: MiR-15a/16 regulates glioma-infiltrating CD8+ T cells activation. International Journal of Cancer. 2017;141: 2082–2092. doi:10.1002/ijc.30912

125. Chu M, Zhao Y, Yu S, Hao Y, Zhang P, Feng Y, et al. miR-15b negatively correlates with lipid metabolism in mammary epithelial cells. Am J Physiol, Cell Physiol. 2018;314: C43–C52. doi:10.1152/ajpcell.00115.2017

